# WRNIP1 PREVENTS G4/R-LOOP-ASSOCIATED GENOMIC INSTABILITY

**DOI:** 10.1101/2022.12.17.520882

**Authors:** Pasquale Valenzisi, Veronica Marabitti, Stefano Petrai, Francesca Rita Giardinelli, Pietro Pichierri, Annapaola Franchitto

**Author notes:** These authors contributed equally to this work (Pasquale Valenzisi, Veronica Marabitti).

## Abstract

Maintenance of genome integrity is essential for cell viability and depends on complete and accurate DNA replication. However, G4/R-loops may provide significant obstacles for DNA replication, as they can cause collisions between replication fork and the transcription machinery. Hence, cells require mechanisms to counteract the presence of persistent G4/R-loops, most of which remain poorly understood. Here, we demonstrate an involvement of the Werner helicase-interacting protein 1 (WRNIP1) in preventing DNA damage induced by G4/R-loop-associated transcription-replication conflicts. We discovered that the ubiquitin-binding domain of WRNIP1 is required to efficiently avoid pathological persistence of G4/R-loops upon replication stress. Also, we observed that G4s reside within R-loops and that WRNIP1 colocalises with these structures. Furthermore, WRNIP1 plays a role in restarting replication from transcription-induced fork stalling. More importantly, we characterized the interplay between WRNIP1 and the DNA helicase FANCJ in counteracting R-loop-dependent G4 formation in response to replication stress. Collectively, our findings propose a mechanisms whereby WRNIP1, contributing to stabilise FANCJ to G4 sites, mitigates the G4/R-loop-mediated transcription-replication conflicts and protects against DNA damage accumulation.

## INTRODUCTION

The genome is continuously threatened by several types of DNA lesions that hinder replication fork progression, impeding the complete and accurate DNA replication prior to cell division. Indeed, protein-DNA complexes, DNA damage, and non-B-form DNA secondary structures can interfere with DNA replication and transcription. Among the variety of non-canonical DNA structures, R-loops have emerged as critical determinants of transcription-replication conflicts (TRCs) that, causing replication stress and genomic instability may promote cancer and other human diseases ^1–3^. R-loops are three-stranded structures, consisting of an RNA-DNA hybrid and a displaced single-stranded DNA, involved in various physiological processes, including transcription termination, regulation of gene expression and DNA repair ^3–5^. Under physiological conditions, R-loops are transient structures thanks to the action of resolving enzymes ^6, 7^. However, if they persist, become pathological provoking the formation of TRCs that, given the high frequency at which the replication and transcription processes take place in cells, are particularly likely to occur. In addition, the interference between transcription and replication at very large human genes may contribute to the instability of common fragile sites (CFS), the most replication stress-sensitive regions in the human genome^8^. Hence, cells have developed a lot of strategies to minimize collisions and help or facilitate replication fork progression that rely on some replication fork protection factors and DNA damage response (DDR) proteins ^9–18^. Recently, an interesting interplay between R-loops and G-quadruplexes-DNA (G4s-DNA) has been observed in cancer cells ^19^. It has been speculated that G-rich sequences in the non-template strand of R-loops can form a G4 motif that play a role in stabilising the R-loop itself ^20^. It is noteworthing that G4s, which are stable DNA secondary structures formed by stacked guanine tetrads held together by Hoogsteen hydrogen bonds that assemble spontaneously on G-rich ssDNA displaced by the moving replication fork, can cause the uncoupling of replication components resulting in fork stalling ^21^. Since these structures are problematic for DNA replication, several DNA helicases deal with G4 resolution to allow a smooth DNA synthesis ^22, 23^. One of them, the FANCJ, a DNA helicase mutated in hereditary breast and ovarian cancer as well as in the chromosomal instability disorder Fanconi anemia, has been recently considered the most potent G4 resolvase ^24^. Therefore, persistent G4/R-loops can promote interference between replication and transcripton machineries, resulting in harmful TRCs. Despite accumulating evidence that, in eukaryotic cells, R-loops and G4s-DNA are a major hindrance for the moving fork and can give rise to replication stress, a hallmark of pre-cancerous and cancerous cells, how exactly cells deal with TRCs caused by persistent G4/R-loops remains largely unknown.

Human WRN-interacting protein 1 (WRNIP1) belongs to the highly conserved AAA+ family of ATPases ^25^. Previous studies revealed that WRNIP1 binds to forked DNA that resembles stalled forks^26^ and its foci overlap with replication factories^27^, suggesting a role at replication forks. Consistent with this, WRNIP1 protects stalled forks from MRE11-mediated degradation and promotes fork restart upon replication stress ^28, 29^. Alongside its role at forks, WRNIP1 is implicated in the activation of the ATM-dependent checkpoint in response to mild replication stress ^30, 31^. More recently, it has been demonstrated that a WRNIP1-mediated response is required to counteract pathological R-loop accumulation in cells with defective ATR-dependent checkpoint activation ^31^. In addition, WRNIP1 was found enriched at CFS, suggesting a role in maintaining their stability ^32^. Interestingly, the best characterised yeast homolog of WRNIP1, Mgs1, binds to G4-forming sites and is required to prevent genomic instability caused by replication arrest ^33, 34^. Nevertheless, a comprehensive understanding of the molecular mechanisms underlying the WRNIP1 function is still missing. Furthermore, it is unclear whether WRNIP1 operates at sites of interference between replication and transcription.

Here, we uncover that the ubiquitin-binding activity of WRNIP1 is required for limiting aberrant accumulation of G4/R-loops that give rise to TRCs. Moreover, we establish that loss of the above WRNIP1 functions is responsible for the elevated genomic instability observed in cells lacking WRNIP1 or its ubiquitin-binding activity in response to mild replication stress. Finally, we demonstrate that WRNIP1 and the DNA helicase FANCJ collaborate in counteracting R-loop-dependent G4 formation upon mild replication stress.

## RESULTS

### Transcription-dependent DNA damage in WRNIP1-deficient and WRNIP1 UBZ mutant cells upon MRS

To investigate the contribution of WRNIP1 in maintaining genome integrity, we evaluated DNA damage accumulation in response to mild replication stress (MRS) induced by nanomolar dose of the DNA polymerase inhibitor aphidicolin (Aph). Beside its ATPase activity, which is involved in fork restart ^28^, WRNIP1 also contains an ubiquitin-binding zinc finger (UBZ) domain that, even if implicated in fork-related functions ^25^, has a still poorly defined function. Hence, in our experiments, we used the SV40-transformed MRC5 fibroblast cell line (MRC5SV), MRC5SV cells stably expressing WRNIP1-targeting shRNA (shWRNIP1) and isogenic cell lines stably expressing the RNAi-resistant full-length wild-type WRNIP1 (shWRNIP1^WT^), its ATPase-dead mutant form (shWRNIP1^T294A^) ^28^ or the UBZ-dead mutant form of WRNIP1 (shWRNIP1^D37A^) that abolishes the ubiquitin-binding activity ^35, 36^ (Figure 1A).

**Figure 1.**
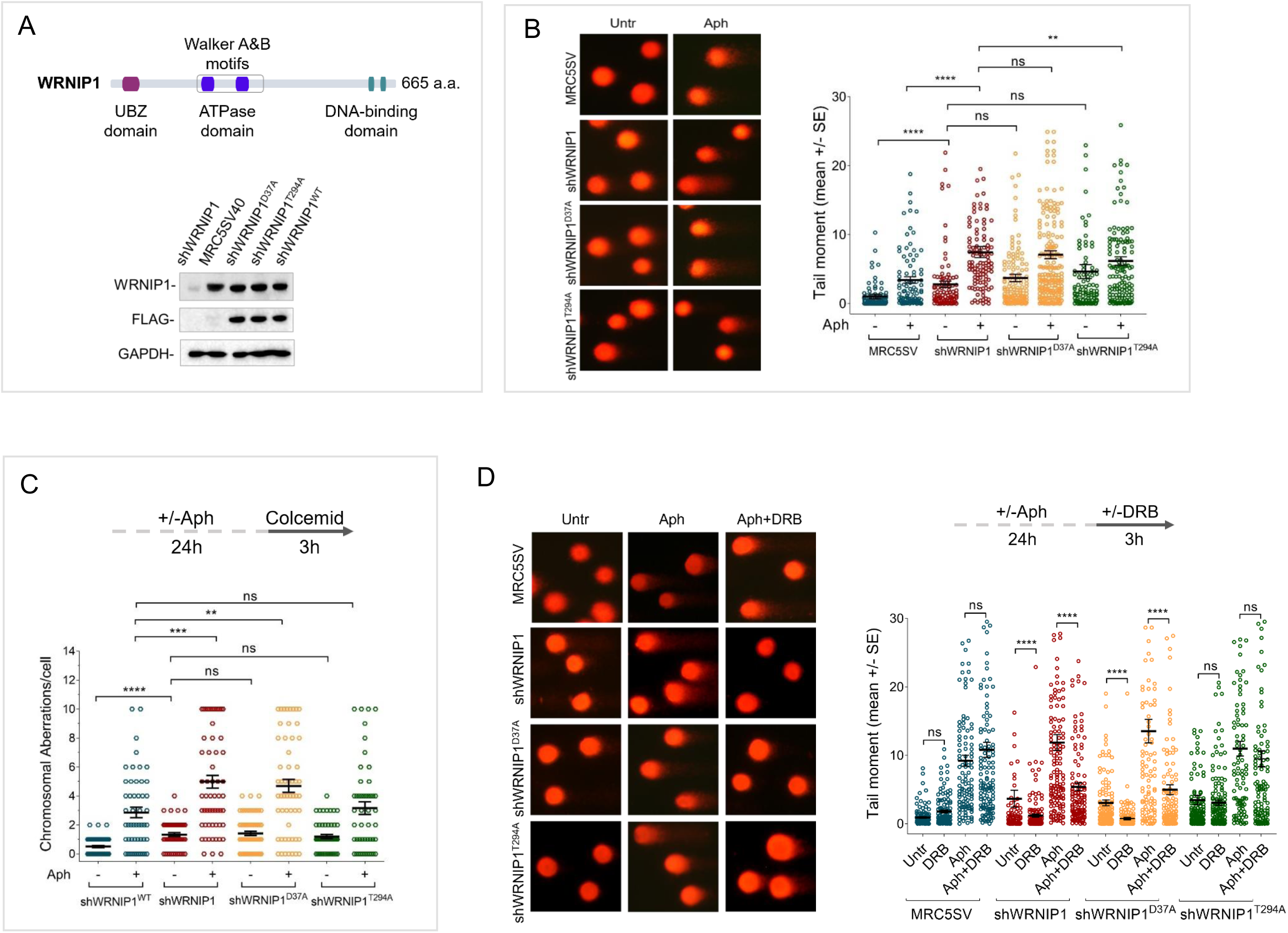
**Loss of WRNIP1 or its UBZ domain results in DNA damage accumulation and enhanced chromosomal instability upon MRS** (**A**) Schematic representation of human WRNIP1 protein structure. Western blot analysis showing the expression of the WRNIP1 protein in wild-type cells (shWRNIP1^WT^), WRNIP1-deficient cells (shWRNIP1) and WRNIP1 ATPase mutant (shWRNIP1^T294A^) or WRNIP1 UBZ mutant (shWRNIP1^D37A^) cells. MRC5SV40 fibroblasts were used as a positive control. The membrane was probed with an anti-FLAG or anti-WRNIP1 antibody. GAPDH was used as a loading control. (**B**) Analysis of DNA damage accumulation evaluated by alkaline Comet assay. MRC5SV, shWRNIP1, shWRNIP1^D37A^ and shWRNIP1^T294A^ cells were treated or not with 0.4 µM Aph for 24 h, and then subjected to Comet assay. The graph shows data presented as mean tail moment ± SE from three independent experiments. Horizontal black lines represent the mean (ns, not significant; **, *P* < 0.01; ****, *P* < 0.0001; two-tailed Student’s t test). Representative images are given. (**C**) Analysis of chromosomal aberrations in the indicated cell lines treated as shown in the experimental scheme. Dot plot shows the number of chromosomal aberrations per cell ± SE from three independent experiments. Horizontal black lines represent the mean (ns, not significant; **, *P* < 0.01; ***, *P* < 0.001; ****, *P* < 0.0001; two-tailed Student’s t test). (**D**) Evaluation of DNA damage accumulation by alkaline Comet assay. Cells were treated as shown in the experimental scheme, and then subjected to Comet assay. Dot plot shows data presented as mean tail moment ± SE from three independent experiments. Horizontal black lines represent the mean (ns, not significant; ****, *P* < 0.0001; two-tailed Student’s t test). Representative images are given.

First, we measured DNA damage by a single-cell electrophoresis in WRNIP1-deficient and WRNIP1 mutant cells. In agreement with previous experiments ^31^, we found that loss of WRNIP1 led to higher spontaneous levels of DNA damage respect to wild-type cells (MRC5SV), and that Aph exacerbated this phenotype (Figure 1B). Notably, in WRNIP1 mutant cells, spontaneous genomic damage was comparable to that observed in WRNIP1-deficient cells but, after MRS, it was significantly enhanced only in WRNIP1 UBZ mutant cells (Figure 1B).

In parallel experiments, we assessed the presence of chromosomal damage. As shown in Figure 1C, shWRNIP1 and WRNIP1 mutant cells exhibited a greater number of chromosomal aberrations under unperturbed conditions, as compared to their corrected isogenic counterparts (shWRNIP1^WT^). However, while loss of WRNIP1 or mutation of its UBZ domain heightened the average number of gaps and breaks after Aph than their corrected counterparts (shWRNIP1^WT^), that of ATPase activity did not (Figure 1C and Suppl Figure 1).

Therefore, loss of WRNIP1 or its ubiquitin-binding function confers high sensitivity of cells to Aph-induced MRS, impacting genomic integrity.

We next wondered whether DNA damage was accumulated in a transcription-dependent manner. To test this possibility, we performed a Comet assay using MRC5SV, shWRNIP1, shWRNIP1^D37A^ and shWRNIP1^T294A^ cells incubated with Aph and/or the 5,6-dichloro-1-ß-D-ribofurosylbenzimidazole (DRB), a strong inhibitor of RNA synthesis, as described ^17^. Our analysis showed that in wild-type and WRNIP1 ATPase mutant cells, DNA damage is not sensitive to transcription inhibition at any of tested conditions (Figure 1D). By contrast, DRB significantly suppressed the amount of DNA damage in both shWRNIP1 and shWRNIP1^D37A^ cells either left untreated or treated with Aph (Figure 1D). Similar results were obtained using another inhibitor of transcription Cordycepin (3’-deoxyadenosine) (^37^; Suppl Figure 2A). Furthermore, anti-phospho-H2AX immunostaining, considered an early sign of DNA damage induced by replication fork stalling ^38^, confirmed a transcription-dependent DNA damage accumulation in shWRNIP1 and shWRNIP1^D37A^ cells (Suppl Figure 2B).

Altogether these findings suggest that WRNIP1 and its UBZ domain are required to suppress transcription-dependent DNA damage accumulation.

### Loss of WRNIP1 or its UBZ domain causes R-loop-dependent DNA damage upon MRS

The main transcription-associated structures that can be detrimental to fork movement, leading to transcription-replication conflicts (TRCs), are R-loops ^39–41^. As transcription-dependent DNA damage accumulation could reflect a role of WRNIP1 in counteracting R-loop-mediated TRCs, we first evaluated the levels of R-loops in our cells, using the well-established anti-RNA-DNA hybrid S9.6 antibody ^17, 42, 43^. A significant increase of the nuclear S9.6 intensity was detected in unperturbed WRNIP1-deficient cells, but not in their corrected counterparts (shWRNIP1^WT^; Figure 2A). Furthermore, although Aph treatment caused an enhancement of R-loop levels in both cell lines, values were considerably more elevated in shWRNIP1 cells (Figure 2A). Notably, removal of R-loops by RNaseH1 overexpression ^44^, strongly suppressed the S9.6 staining (Figure 2A). To further confirm R-loop accumulation within nuclear DNA, we isolated genomic DNA from shWRNIP1^WT^ and shWRNIP1 cells, and performed dot blot analysis. Consistent with fluorescence analysis, the S9.6 signal was higher in shWRNIP1 cells than in shWRNIP1^WT^ cells and was abolished by RNaseH treatment (Figure 2B; ^45^). This result raises the possibility that DNA damage accumulation observed in WRNIP1-deficient cells is correlated to the elevated accumulation of R-loops.

**Figure 2.**
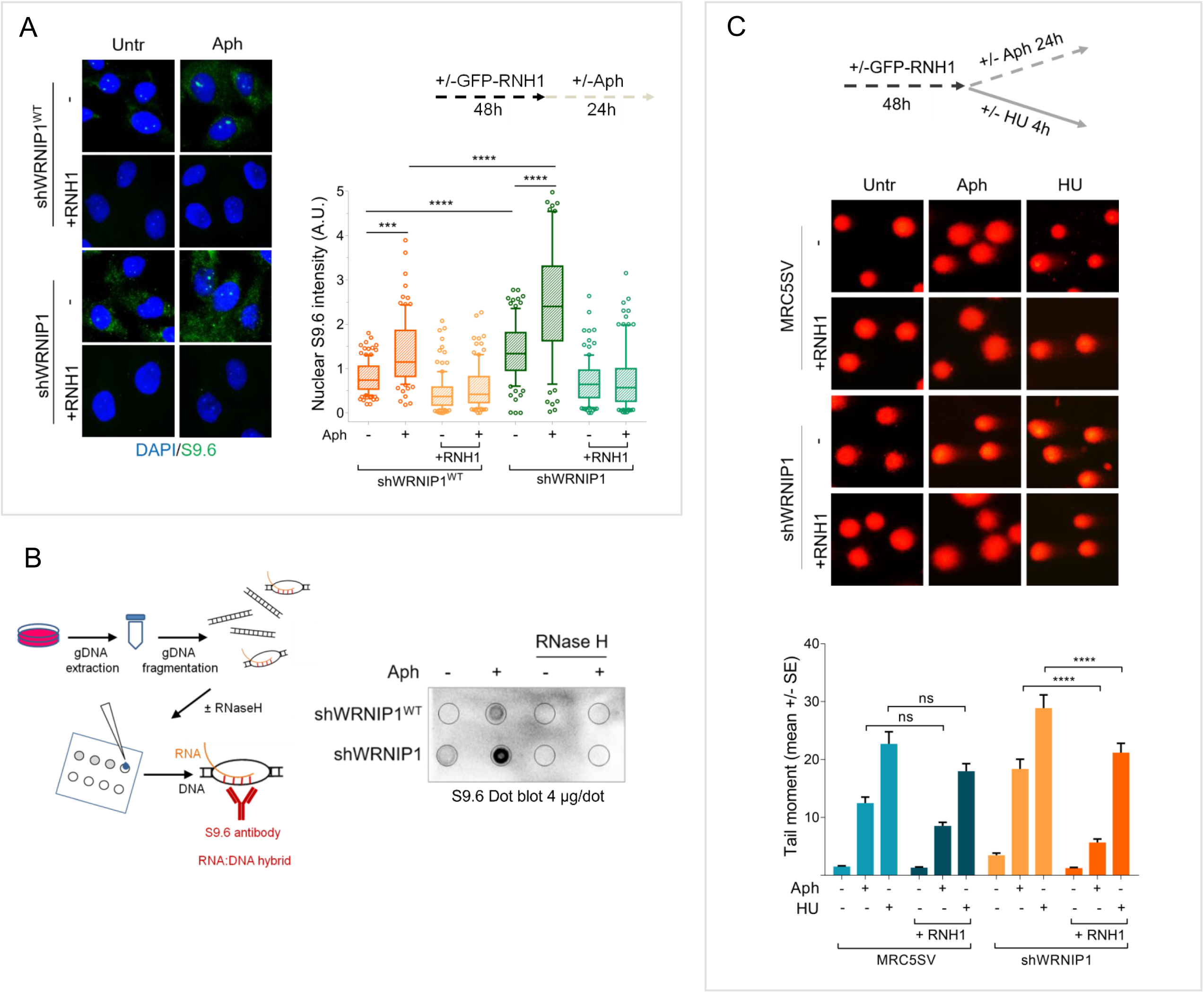
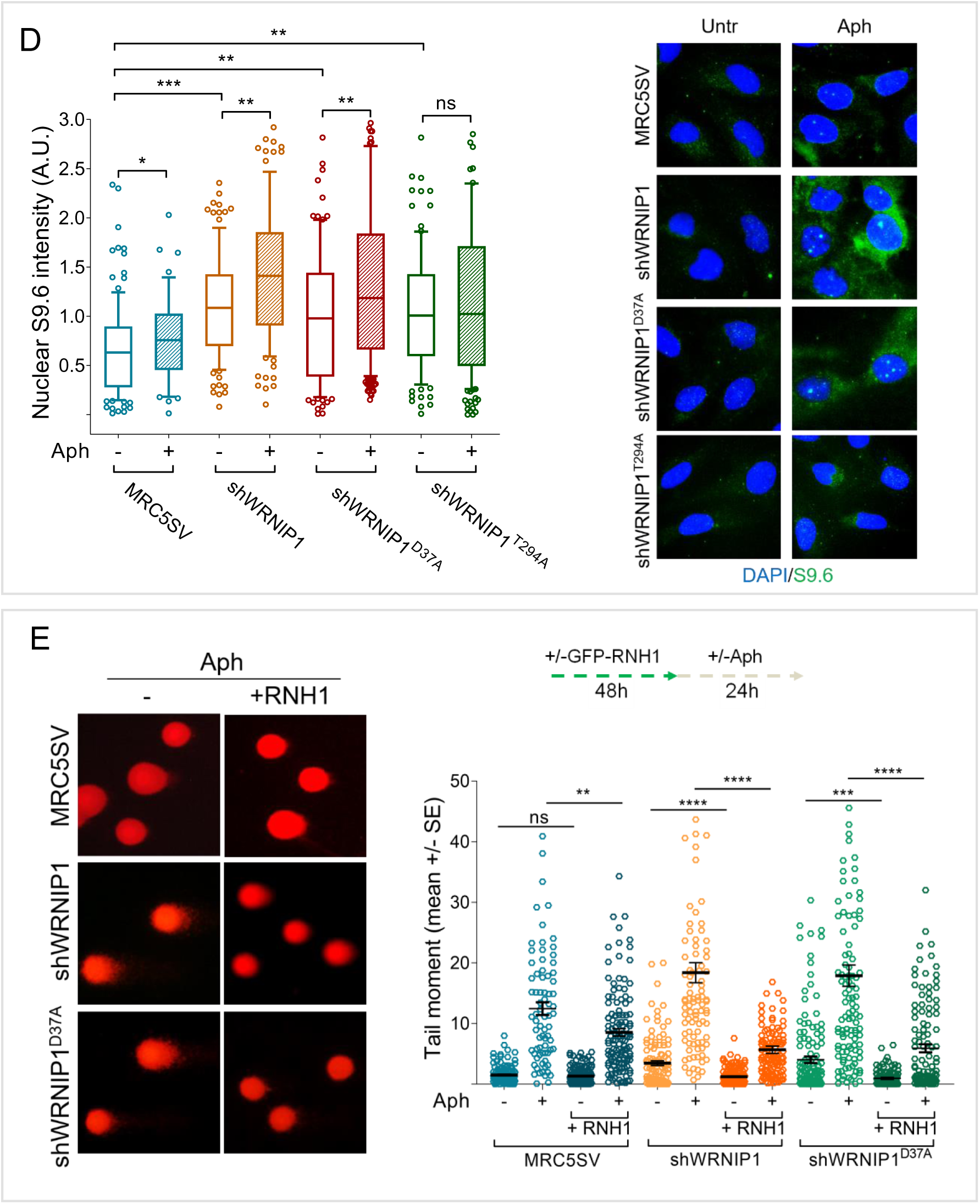
**Loss of WRNIP1 or its UBZ domain results in R-loop-dependent accumulation upon MRS** (**A**) Evaluation of R-loop accumulation by immunofluorescence analysis in shWRNIP1^WT^ and shWRNIP1 cells treated as reported in the experimental design after transfection with GFP-tagged RNaseH1 or empty vector. Cells were fixed and stained with anti-RNA-DNA hybrid S9.6 monoclonal antibody. Representative images are given. Nuclei were counterstained with DAPI. Box plot shows nuclear S9.6 fluorescence intensity. Box and whiskers represent 20-75 and 10-90 percentiles, respectively. The line represents the median value. Data are presented as means of three independent experiments. Horizontal black lines represent the mean. Error bars represent standard error (***, *P* < 0.001; ****, *P* < 0.0001; two-tailed Student’s t test). (**B**) Dot blot to confirm R-loop accumulation. Genomic DNA isolated from shWRNIP1WT and shWRNIP1 cells, treated as reported in the experimental design and processed as described in “Materials and Methods”, was spotted onto nitrocellulose membrane. The membrane was probed with anti-RNA-DNA hybrid S9.6 monoclonal antibody. Treatment with RNase H was used as a negative control. Representative gel images of at least three replicates are shown. (**C**) Analysis of DNA damage accumulation by alkaline Comet assay. MRC5SV and shWRNIP1 cells were treated or not with Aph or HU, as reported by the experimental scheme after transfection with GFP-tagged RNaseH1 or empty vector (-), and then subjected to Comet assay. The graph shows data presented as mean tail moment ± SE from three independent experiments (ns, not significant; **** *P* < 0.0001; Mann-Whitney test). Representative images are given. (**D**) Immunofluorescence analysis to determine R-loop levels in MRC5SV, shWRNIP1, shWRNIP1^D37A^ and shWRNIP1^T294A^ cells treated or not with 0.4 µM Aph for 24 h. Cells were fixed and stained with anti-RNA-DNA hybrid S9.6 monoclonal antibody. Representative images are given. Nuclei were counterstained with DAPI. Box plot shows nuclear S9.6 fluorescence intensity. Box and whiskers represent 20-75 and 10-90 percentiles, respectively. The line represents the median value. Data are presented as means of three independent experiments. Horizontal black lines represent the mean. Error bars represent SE (ns, not significant; * *P* < 0.05; ** *P* < 0.01; ***, *P* < 0.001; two-tailed Student’s t test). (**E**) Analysis of the effect of R-loop resolution on DNA damage accumulation evaluated by alkaline Comet assay. Cells were treated as reported in the experimental scheme after transfection with GFP-tagged RNaseH1 or empty vector, and then subjected to Comet assay. Dot plot shows data presented as mean tail moment ± SE from three independent experiments. Horizontal black lines represent the mean (ns, not significant; ** *P* < 0.01; *** *P* < 0.001; **** *P* < 0.0001; Mann-Whitney test). Representative images are given.

Having demonstrated that WRNIP1-deficient cells accumulate high levels of R-loops, we explored the risks for genome integrity comparing different replication stress conditions. As observed above, loss of WRNIP1 slightly increased the amount of spontaneous DNA damage, while Aph enhanced comet tail moment with values greater in shWRNIP1 than in MRC5SV cells (Figure 2C). A similar trend was obtained using hydroxyurea (HU) as replication-perturbing agent (Figure 2C). It is noteworthy that RNaseH1 overexpression was able to strongly reduce tail moment only in shWRNIP1 cells, and more efficiently after Aph-induced replication slowdown than arrest by HU (Figure 2C), confirming R-loops as a driver of genome instability in WRNIP1-deficient cells and suggesting that R-loop-mediated DNA damage is more evident after a replication stress that does not block completely replication fork progression.

Next, we asked which of the WRNIP1 activities could prevent formation of aberrant R-loops upon MRS. A similar level of spontaneous R-loops was observed in cells with mutations in either ATPase or UBZ domain. However, after MRS, shWRNIP1^D37A^ cells exhibited a more pronounced S9.6 signal than shWRNIP1^T294A^ cells (Figure 2D). This result reflects a role of UBZ domain of WRNIP1 in counteracting R-loop accumulation upon MRS.

Finally, we verified whether R-loop-mediated DNA damage in WRNIP1-deficient cells could depend on loss of UBZ domain of WRNIP1. Importantly, we noted that suppression of DNA damage in shWRNIP1^D37A^ cells upon RNaseH1 overexpression was similar to that seen in shWRNIP1 cells (Figure 2E), suggesting that ubiquitin-binding activity of WRNIP1 is needed to avoid an R-loop-mediated DNA damage upon MRS.

### UBZ domain of WRNIP1 is required to attenuate TRCs upon MRS

Since TRCs play a crucial role in promoting R-loop-mediated genomic instability ^46^, we investigated the occurrence of such conflicts in our conditions. Hence, we performed an established proximity ligation assay (PLA) that allows to detect physical interactions ^47^ by using antibodies against proliferating cell nuclear antigen (PCNA) and RNA polymerase II (RNA pol II) to mark replication forks and transcription complexes, respectively, as previously reported ^43^. Our analysis showed greater spontaneous PLA signals (red spots) in WRNIP1-deficient and WRNIP1 UBZ mutant cells than in wild-type cells (Figure 3A). Notably, although Aph increased the co-localization of PCNA and RNA pol II in all cell lines, however the number of PLA spots was significantly higher in shWRNIP1 and shWRNIP1^D37A^ cells than in shWRNIP1^WT^ cells (Figure 3A). Interestingly, this phenotype is considerably diminished by RNaseH1 overexpression (Figure 3A), suggesting that the UBZ domain of WRNIP1 can play a role in attenuating R-loop-induced TRCs upon MRS.

**Figure 3.**
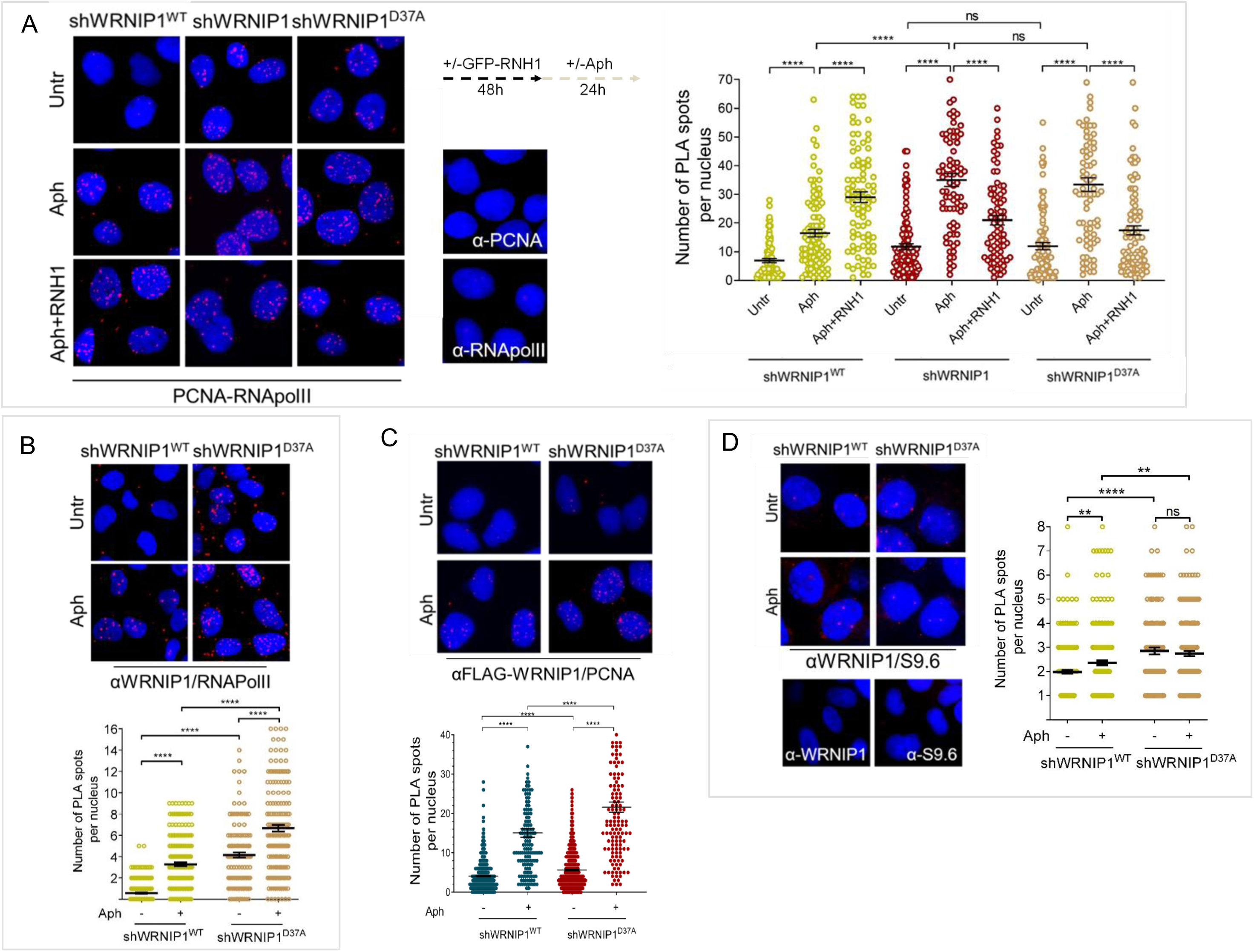
**Loss of WRNIP1 or its UBZ domain promotes R-loop-dependent TRCs accumulation** (**A**) Detection of TRCs by fluorescence-based PLA assay in MRC5SV, shWRNIP1 and shWRNIP1^D37A^ cells treated as reported in the experimental scheme after transfection with GFP-tagged RNaseH1 or empty vector. Cells were fixed and stained with antibodies against PCNA and RNA pol II. Representative images are given. Each red spot represents a single interaction between proteins. No spot has been revealed in cells stained with each single antibody (negative control). Nuclei were counterstained with DAPI. Dot plot shows the number of PLA spots per nucleus. Data are presented as means of three independent experiments. Horizontal black lines represent the mean ± SE (ns, not significant; ****, *P* < 0.0001; Mann-Whitney test). (**B** and **C**) Analysis of localization of WRNIP1 near/at the transcription and replication machineries. Cells were treated or not with 0.4 µM Aph for 24 h, fixed and stained with antibodies against WRNIP1 and RNA pol II (**B**) or WRNIP1 and PCNA (**C**) to visualize the interaction between WRNIP1 and replication or transcription machinery, respectively. Each red spot represents a single interaction between proteins. Representative images are given. Nuclei were counterstained with DAPI. Dot plots show the number of PLA spots per nucleus. Data are presented as means of three independent experiments. Horizontal black lines represent the mean ± SE (****, *P* < 0.0001; Mann-Whitney test). (**D**) Detection of physical interaction between WRNIP1 and R-loops. Cells were treated or not with 0.4 µM Aph for 24 h, and then subjected to PLA assay as described in “Supplementary Materials and Methods”. Cells were stained with anti-RNA-DNA hybrid S9.6 antibody, and an antibody raised against WRNIP1. Representative images are given. Each red spot represents a single interaction between R-loops (S9.6) and WRNIP1. No spot has been revealed in cells stained with each single antibody (negative control). Nuclei were counterstained with DAPI. Dot plot shows the number of PLA spots per nucleus. Horizontal black lines represent the mean ± SE (** *P* < 0.01****, *P* < 0.0001; Mann-Whitney test).

We next examined the localization of WRNIP1 near/at the transcription and replication machineries, performing PLA assays in shWRNIP1^WT^ and shWRNIP1^D37A^ cells using antibodies against WRNIP1 and RNA pol II or PCNA, respectively. Consistent with the possibility that WRNIP1 localizes at the sites of TRCs, we observed increased numbers of PLA spots between WRNIP1 and RNA pol II (Figure 3B) or WRNIP1 and PCNA (Figure 3C) upon Aph in wild-type cells. Surprisingly, a phenotype similar but more evident was observed in WRNIP1 UBZ mutant cells respect to wild-type cells, under both unperturbed and treatment conditions (Figures 3B and C).

Using PLA assay, we also explored the ability of WRNIP1 to localise at/near R-loops, and we noted an increasing number of interactions between WRNIP1 and R-loops (anti-S9.6) (Figure 3D), supporting the hypothesis of a role of WRNIP1 in response to R-loop accumulation.

Taken together, these findings suggest an increased co-localization of WRNIP1 with transcription/replication complexes and R-loops following MRS and implicate ubiquitin-binding function of WRNIP1 in mitigating R-loop-induced TRCs.

### Transcription-mediated R-loop formation is a barrier to DNA replication in cells lacking WRNIP1 or its UBZ domain

R-loops are the main transcription-associated structures that have the capacity to hinder replication fork progression ^48^ . To directly evaluate the impact of R-loops on fork progression, we performed DNA fiber assay and examined replication fork dynamics at single molecule resolution in Aph-treated MRC5SV, shWRNIP1 and WRNIP1 UBZ mutant cells. Cells were sequentially labelled with the thymidine analogues 5-chloro-2’-deoxyuridine (CldU) and 5-iodo-2’-deoxyuridine (IdU) as described in the scheme (Figure 4A). Under normal growth conditions, and in agreement with previous data ^28^, MRC5SV and shWRNIP1 cells showed almost similar fork velocity, while it was significantly reduced in shWRNIP1^D37A^ cells (Figure 4A). After Aph treatment, although fork velocity decreased in all cell lines, values were considerably lower when WRNIP1 or its UBZ domain was lost (Figure 4B). Importantly, RNaseH1 overexpression led to a significant enhancement in the fork progression rate only in shWRNIP1 and shWRNIP1^D37A^ cells (Figure 4A and B). DNA fiber analysis also showed that loss of WRNIP1 or its UBZ domain resulted in a greater percentage of stalled forks induced by Aph with respect to control cells (Figure 4C). Furthermore, comparing the percentage of restarting forks in all cell lines, we noted that the absence of WRNIP1 reduced the ability of cells to resume replication after release from Aph in the same extent as loss of its UBZ domain (Figure 4D). Suppressing R-loop formation, the percentage of stalled forks was lowered and, consistently, that of restarting forks augmented in both shWRNIP1 and shWRNIP1^D37A^ cells (Figure 4 C and D). Similar results were obtained treating cells with DRB (Suppl. Figure 3A-D).

**Figure 4.**
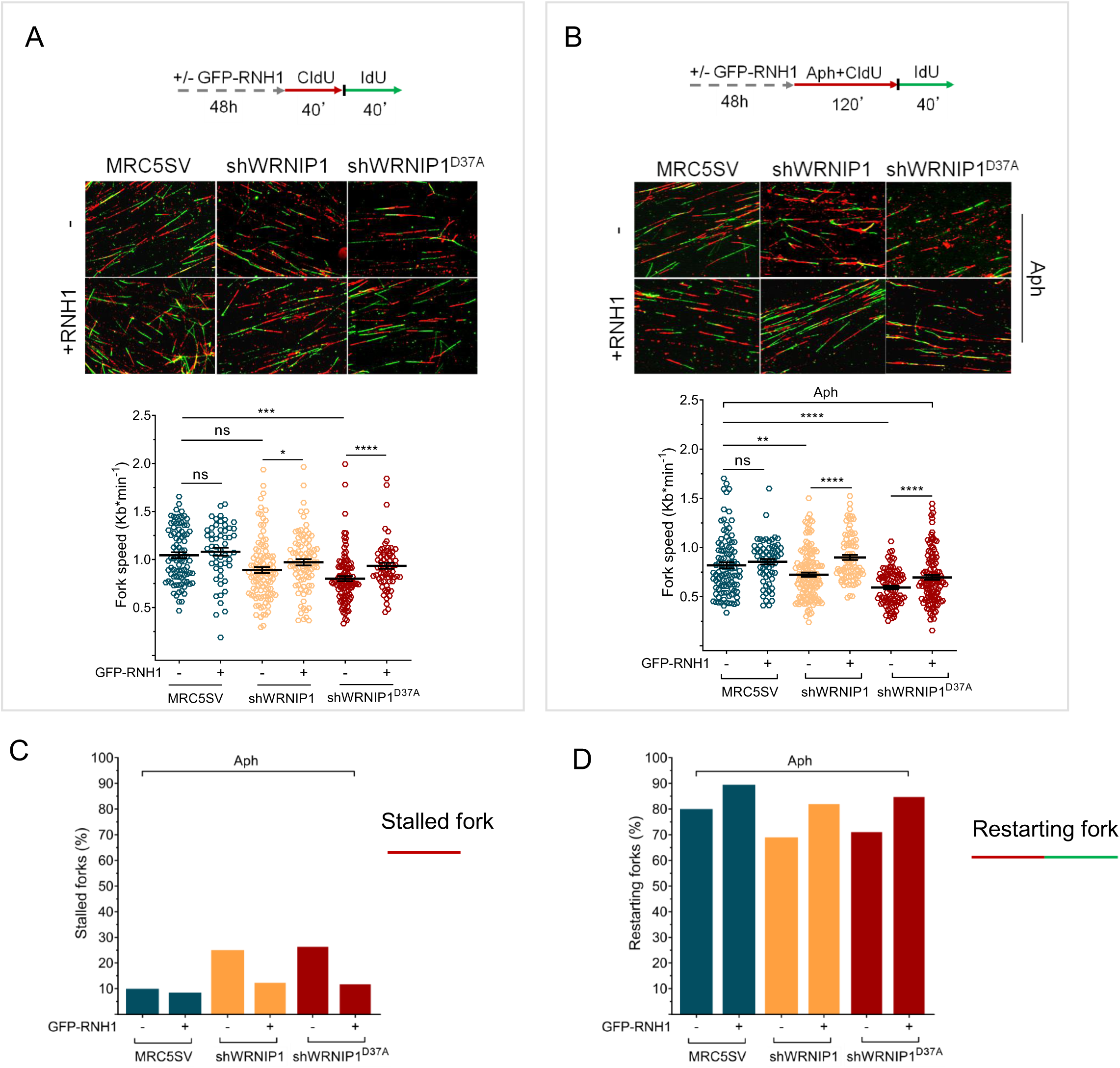
**R-loops affects DNA replication in cells lacking WRNIP1 or its UBZ domain upon MRS** Experimental scheme of dual labelling of DNA fibers in MRC5SV, shWRNIP1 and shWRNIP1^D37A^ cells under unperturbed conditions (**A**) or upon MRS (**B**). After transfection with GFP-tagged RNaseH1 or empty vector (-), cells were pulse-labelled with CldU, treated or not with 0.4 µM Aph and then subjected to a pulse-labelling with IdU. Representative DNA fiber images are shown. The graph shows the analysis of replication fork velocity (fork speed) in the cells. The length of the green tracks was measured. Mean values are represented as horizontal black lines (ns, not significant; *, *P* < 0.05; **, *P* < 0.01; ***, *P* < 0.001; ****, *P* < 0.0001; Mann-Whitney test). (**C** and **D**) The graph shows the percentage of red (CldU) tracts (stalled forks) or green (IdU) tracts (restarting forks) in the cells. Error bars represent standard error (ns, not significant; *, *P* < 0.05; Mann-Whitney test).

Therefore, we concluded that R-loops provide a significant obstacle for replication in cells lacking WRNIP1 or its UBZ domain, and that ubiquitin-binding activity of WRNIP1 can play a crucial function in restarting replication from transcription-induced fork stalling.

### Loss of WRNIP1 or its UBZ domain increases G4/R-loop levels

It has been reported that G4 formation is strongly correlated with the presence of R-loops, which are stabilized by G4s ^19, 20^. Hence, we assumed that G4/R-loops are formed in the absence of WRNIP1 or its UBZ domain and that, creating an obstacle to moving fork, they can lead to enhanced TRCs in response to replication stress. To test this hypothesis, we first evaluated the amount of G4s in cells treated or not with Aph, employing a well-established G-quadruplex structure-specific antibody ^49^.

A pronounced spontaneous increase in BG4 nuclear intensity was detected in WRNIP1-deficient and WRNIP1 UBZ mutant cells, but not in WRNIP1 ATPase mutant cells and the corrected counterparts (Figure 5A). Notably, Aph caused a significant enrichment of G4 levels only in shWRNIP1 and shWRNIP1^D37A^ cells respect to wild-type cells, and transcription inhibition relieved this phenotype (Figure 5A). Consistently, using pyridostatin (PDS), a G4-stabilizing ligand ^50^, although the BG4 signal was enhanced in all cell lines, a significant G4 accumulation was detected exclusively in the absence of WRNIP1 or its ubiquitin-binding domain when compared to wild-type cells (Suppl. Figure 4A). Importantly, inhibiting ubiquitin ligase RAD18, which monoubiquitinates PCNA at stalled replication forks with which WRNIP1 preferentially interacts ^30^, enhanced interaction between WRNIP1 and G4s (Suppl. Figure 4B).

**Figure 5.**
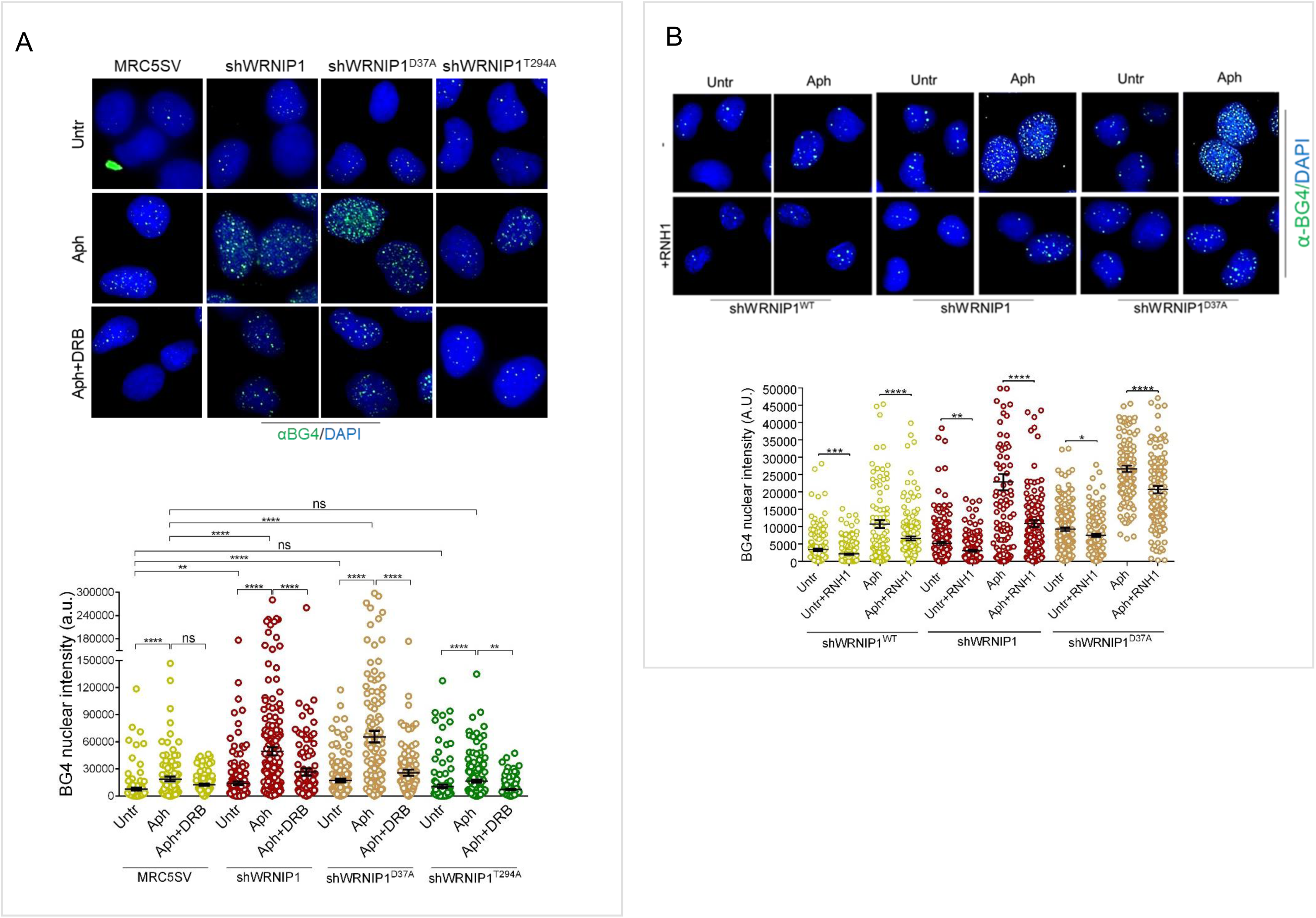
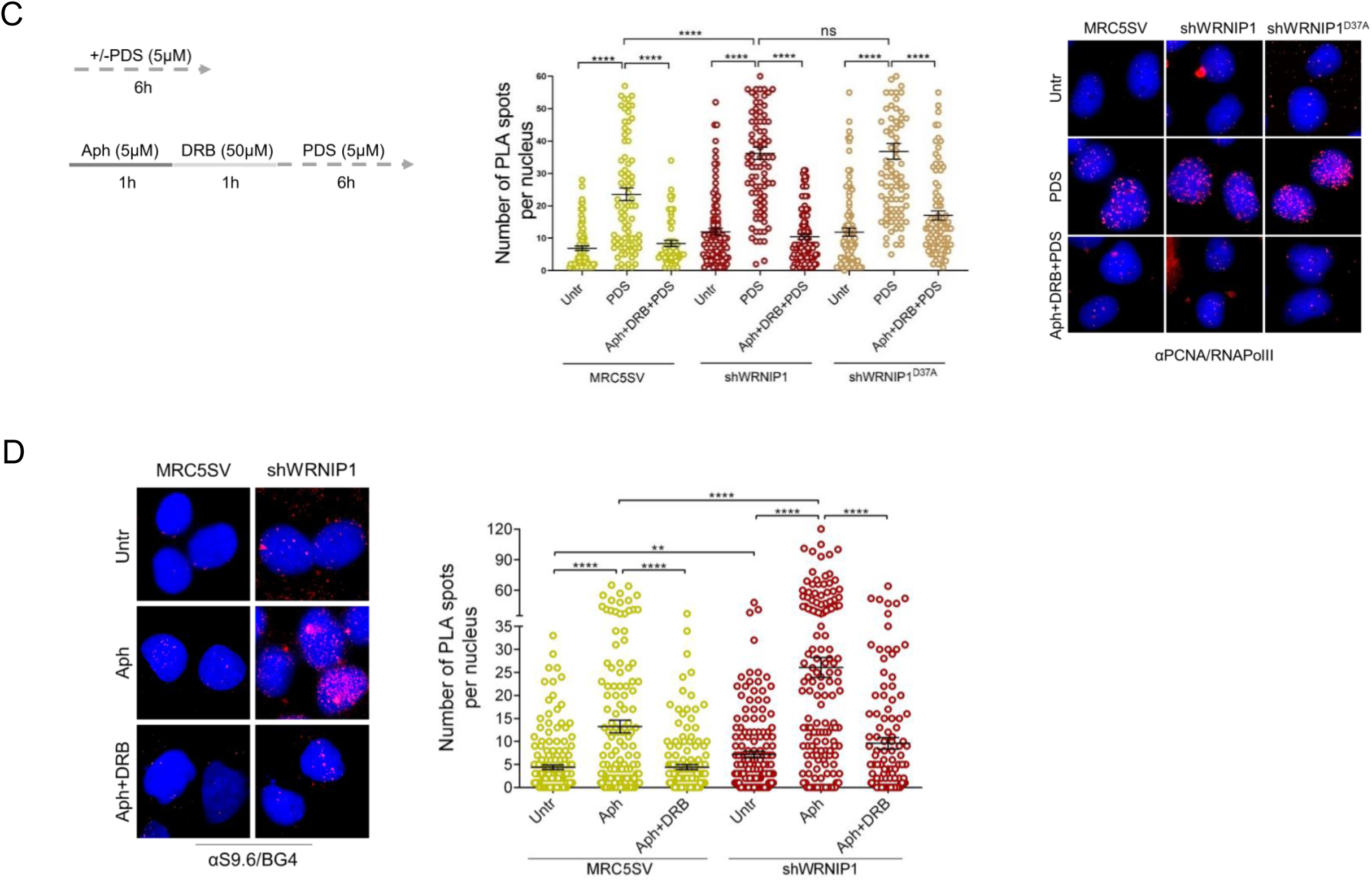
**G4/R-loops causes TCRs in cells lacking WRNIP1 or its UBZ domain** (**A**) Evaluation of G4 levels by immunofluorescence of shWRNIP1^WT^, shWRNIP1, shWRNIP1^D37A^ and shWRNIP1^T294A^ cells treated or not with 0.4 µM Aph for 24 h and the transcription inhibitor DRB the last 3 h. Visualization of DNA G4s was performed with the BG4 antibody as described in “Materials and Methods”. Immunostaining shows G4s foci (green) in the cells. Nuclei were counterstained with DAPI. Representative images are given. The graph shows the quantification of BG4 nuclear intensity per nucleus. Data are presented as means of three independent experiments. Horizontal black lines represent the mean ± SE (ns, not significant; ****, *P* < 0.0001; Mann-Whitney test). (**B**) Immunostaining with BG4 in shWRNIP1^WT^and shWRNIP1 cells treated or not with 0.4 µM Aph for 24 h after transfection with GFP-tagged RNaseH1 or empty vector. Visualization of G4s was carried out as in (A). Representative images are given. The graph shows the quantification of BG4 nuclear intensity per nucleus. Data are presented as means of three independent experiments. Horizontal black lines represent the mean ± SE (ns, not significant; ** *P* < 0.01; ***, *P* < 0.001; ****, *P* < 0.0001; Mann-Whitney test). (**C**) PLA assay to assess TRCs in MRC5SV, shWRNIP1 and shWRNIP1^D37A^ cells treated as reported in the experimental scheme. Cells were fixed and stained with antibodies against PCNA and RNA pol II. Representative images are given. Each red spot represents a single interaction between proteins. Nuclei were counterstained with DAPI. Dot plot shows the number of PLA spots per nucleus. Data are presented as means of three independent experiments. Horizontal black lines represent the mean ± SE (ns, not significant; ****, *P* < 0.0001; Mann-Whitney test). (**D**) Analysis of the co-localization of G4 and R-loop by fluorescence-based PLA assay in MRC5SV and shWRNIP1 treated or not with 0.4 µM Aph for 24 h alone or in combination with DRB 50 µM for the last three hours. Cells were fixed and stained with BG4 and S9.6 antibodies. Representative images are given. Each red spot represents a single interaction between G4 and R-loop. Nuclei were counterstained with DAPI. Dot plot shows the number of PLA spots per nucleus. Data are presented as means of three independent experiments. Horizontal black lines represent the mean ± SE (** *P* < 0.01; ****, *P* < 0.0001; Mann-Whitney test).

We therefore considered the possibility that enhanced BG4 signal observed in shWRNIP1 and shWRNIP1^D37A^ cells upon Aph could depend on G4s residing within R-loops. In accordance with this, we observed that RNaseH1 overexpression strongly reduced BG4 nuclear intensity (Figure 5B). In agreement with the observation that G4s stabilisation of co-transcriptional R-loops can promote TRCs, PDS improved such conflicts in shWRNIP1 and shWRNIP1^D37A^ cells at higher levels in comparison with shWRNIP1^WT^ cells and are prevented by pre-treatment with replication (Aph) and transcription (DRB) inhibitors ^51^, reaching values of untreated samples (Figure 5C). Furthermore, PDS-induced G4 formation in cells was precluded by preventing TRCs (Suppl. Figure 5A).

These results suggest that loss of WRNIP1 or its UBZ domain results in a robust TRCs-dependent accumulation of G4 structures.

Next, enhanced co-localization of G4s and R-loops was visualized in shWRNIP1 cells and repressed by DRB treatment (Figure 5D). To further strengthen an involvement of WRNIP1 in G4/R-loops formation and given that TRCs can arise because of a failure to resolve these structures, we assessed R-loop formation upon short or prolonged exposure to PDS. We found that nuclear R-loops were strengthened by G4 stabilization, in agreement with a previous study ^19^, and that levels were significantly elevated in the absence of WRNIP1 or its UBZ domain (Suppl. Figure 5B and Figure 6A).

**Figure 6.**
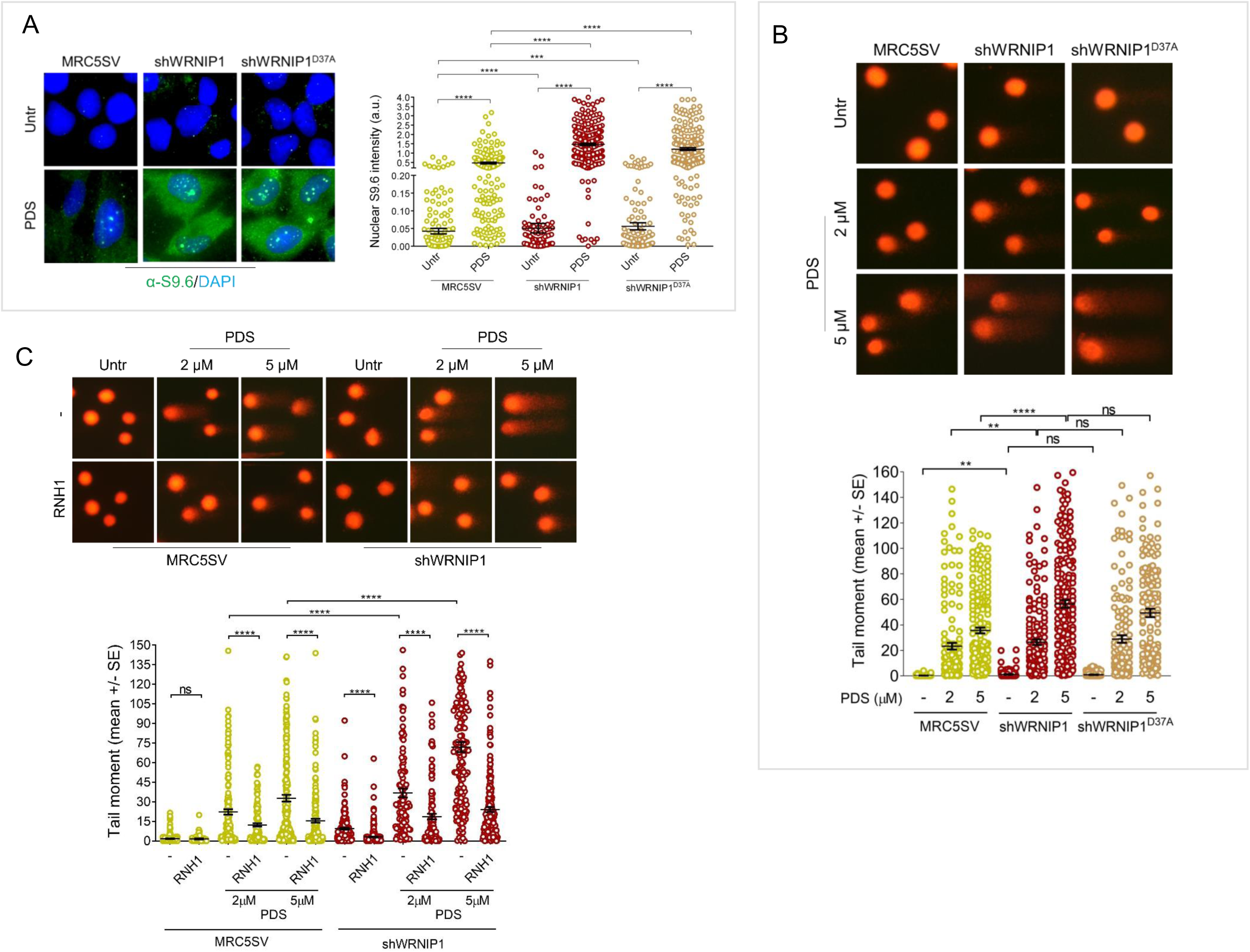
**G4/R-loop-dependent DNA damage in cells lacking WRNIP1 or its UBZ domain** (**A**) Evaluation of R-loop levels by immunofluorescence analysis in MRC5SV, shWRNIP1 and shWRNIP1^D37A^ cells treated or not with 5 µM PDS for 24 h. Cells were fixed and stained with anti-RNA-DNA hybrid S9.6 monoclonal antibody. Representative images are given. Nuclei were counterstained with DAPI. The graph shows nuclear S9.6 fluorescence intensity. The line represents the median value. Data are presented as means of three independent experiments. Horizontal black lines represent the mean. Error bars represent standard error (***, *P* < 0.001; ****, *P* < 0.0001; Mann-Whitney test). (**B**) Analysis of DNA damage accumulation by alkaline Comet assay. MRC5SV and shWRNIP1 cells were treated or not with 2 or 5 µM PDS for 24 h, and then subjected to Comet assay. The graph shows data presented as mean tail moment ± SE from three independent experiments (ns, not significant; **, *P* < 0.01; **** *P* < 0.0001; Mann-Whitney test). Representative images are given. (**C**) Effect of R-loop resolution on DNA damage accumulation evaluated by alkaline Comet assay in cells were treated or not with 2 or 5 µM PDS for 24 h after transfection with GFP-tagged RNaseH1 or empty vector. Dot plot shows data presented as mean tail moment ± SE from three independent experiments. Horizontal black lines represent the mean (ns, not significant; **** *P* < 0.0001; Mann-Whitney test). Representative images are given.

Finally, we asked whether PDS induced DNA damage due to the presence of R-loops in our cells. Alkaline Comet assay revealed that loss of WRNIP1 or its UBZ domain caused higher spontaneous levels of DNA damage respect to wild-type cells (MRC5SV), and that PDS exacerbated this phenotype in a dose-dependent manner (Figure 6B). In line with previous data ^51^, neutral Comet assay confirmed the presence of double-strand breaks (DSBs) in WRNIP1-deficient and WRNIP1 UBZ mutant cells treated with PDS (Suppl. Figure 6A). More interestingly, overexpression of RNaseH1 markedly lowered both DNA damage and DSB formation after PDS in the absence of WRNIP1 (Figure 6C and Suppl. Figure 6B). Also, counteracting TRCs restricted DSB formation in WRNIP1-deficient cells (Suppl. Figure 7).

Taken together, these findings show that G4/R-loops, responsible for the onset of TCRs, arise due to loss of WRNIP1 or its UBZ domain and leads to DNA damage.

### WRNIP1 loss increased G4-ssDNA interaction upon MRS

Recently, an interesting interplay between R-loops and G4s has been observed in cancer cells ^19^. It has been speculated that G-rich sequences in the non-template strand of R-loops can form a G4 motif that play a role in stabilising the R-loop itself ^20^. First, we decided to evaluate the levels of ssDNA in the absence of WRNIP1 upon Aph-or PDS-induced replication stress. To this aim, MRC5SV and shWRNIP1 cells were pre-labelled with the thymidine analogue 5-iodo-2’-deoxyuridine (IdU) and treated with Aph for different times as described in the scheme (Figure 7A). We specifically visualized ssDNA formation at parental-strand by immunofluorescence using an anti-BrdU/IdU antibody under non-denaturing conditions as reported ^52^. Our analysis showed that WRNIP1-deficient cells presented a significant higher amount of ssDNA than wild-type cells upon Aph treatment (Figure 7A). Notably, transcription inhibition caused a significant reduction of ssDNA levels in shWRNIP1 cells (Figure 7A). As DRB suppresses the G4 levels upon Aph treatment in shWRNIP1 cells (Figure 6A), it is likely that the amount of ssDNA accumulated in the absence of WRNIP1 can be attributable mainly to the detection of the non-template strand of R-loops where G4s can form.

**Figure 7.**
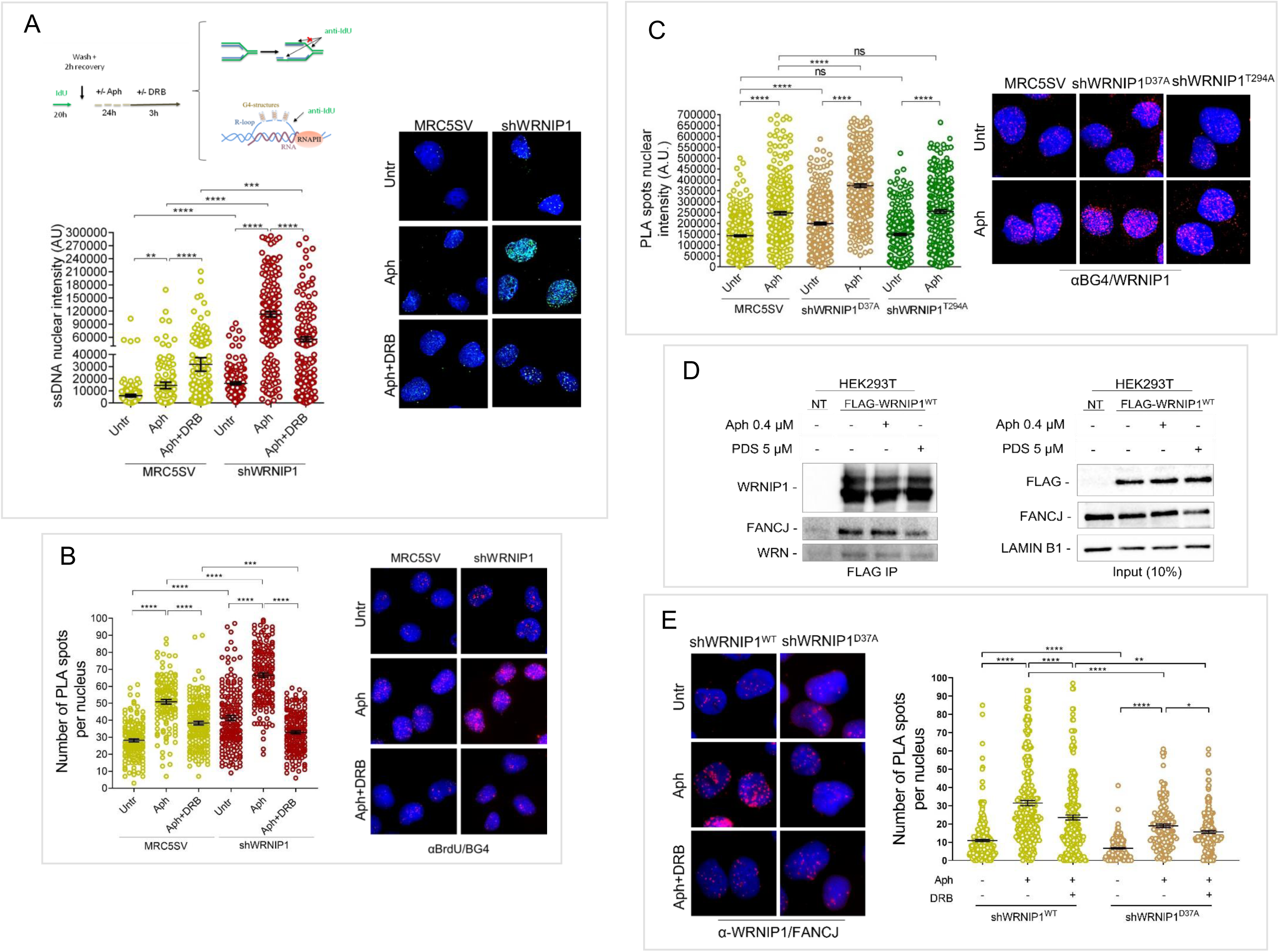
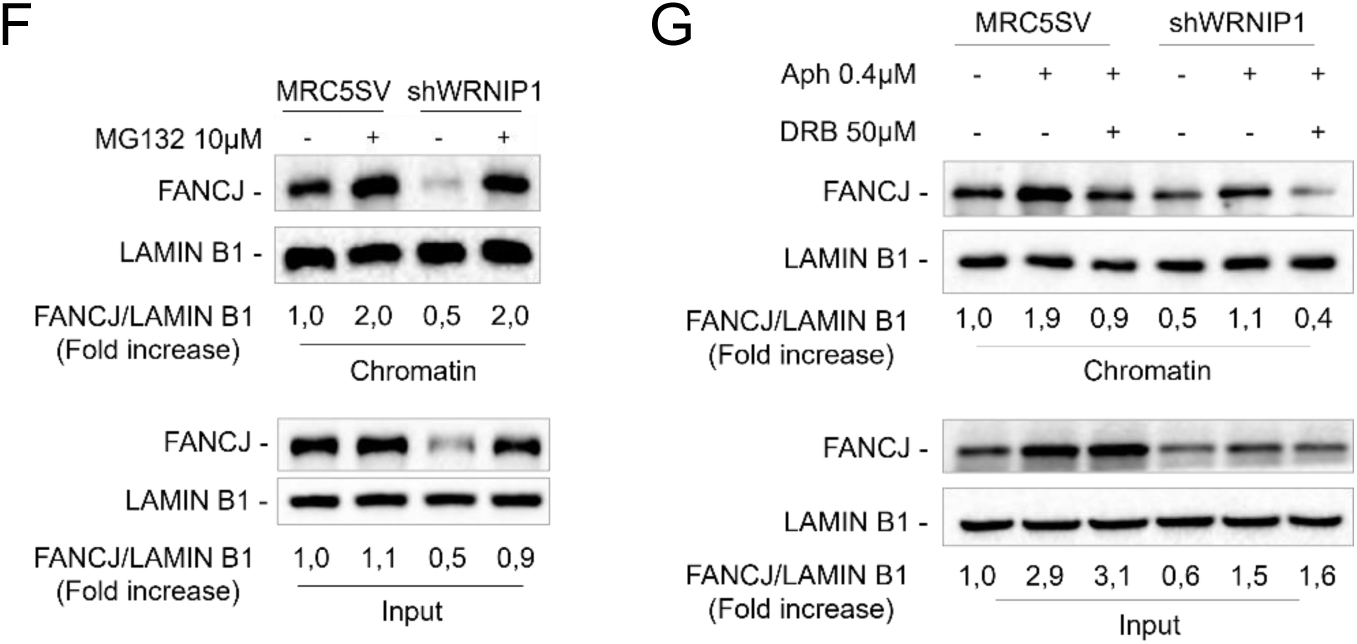
**Analysis of the interaction between WRNIP1 and FANCJ in response to G4/R-loop accumulation** (**A**) Evaluation of parental ssDNA exposure by immunofluorescence analysis in MRC5SV and shWRNIP1 cells. Cells were treated as described in the experimental scheme and then fixed and stained with anti-IdU antibody. Representative images are given. Nuclei were counterstained with DAPI. The graph shows nuclear ssDNA fluorescence intensity. Data are presented as means of three independent experiments. Horizontal black lines represent the mean. Error bars represent standard error (**, *P* < 0.01; ****, *P* < 0.0001; Mann-Whitney test). (**B**) Analysis of the co-localization of G4s and parental ssDNA by PLA assay. Cells were treated as in (A) and stained with anti-BrdU and BG4 antibodies. Representative images are given. Each red spot represents a single interaction between ssDNA and G4s. Nuclei were counterstained with DAPI. Dot plot shows the number of PLA spots per nucleus. Data are presented as means of three independent experiments. Horizontal black lines represent the mean ± SE (*** *P* < 0.001 ****, *P* < 0.0001; Mann-Whitney test). (**C**) PLA assay to explore the colocalization of WRNIP1 with G4-structures in shWRNIP1^WT^ and shWRNIP1^T294A^ cells with or without 0.4 µM Aph treatment for 24 h. Cells were fixed and stained with anti-WRNIP1 and BG4 antibodies. Representative images are given. Each red spot represents a single interaction between WRNIP1 and G-quadruplex. Nuclei were counterstained with DAPI. Dot plot shows the number of PLA spots per nucleus. Data are presented as means of three independent experiments. Horizontal black lines represent the mean ± SE (ns, not significant; ****, *P* < 0.0001; Mann-Whitney test). (**D**) Co-IP analysis showing the interaction between WRNIP1 and FANCJ after IP with anti-FLAG antibody in HEK293T cell extracts treated or not with 0.4 µM Aph or 5 µM PDS for 24 h, followed by WB. The membranes were probed with the indicated antibodies. LAMIN B1 was used as a loading control. (**E**) Analysis of the interaction between WRNIP1 and FANCJ in MRC5SV and shWRNIP1^D37A^ cells treated or not with 0.4 µM Aph for 24 h and the transcription inhibitor DRB the last 3 h. Representative images are given. Each red spot represents a single interaction between WRNIP1 and FANCJ. Nuclei were counterstained with DAPI. Dot plot shows the number of PLA spots per nucleus. Data are presented as means of three independent experiments. Horizontal black lines represent the mean ± SE (ns, not significant; * *P* < 0.05; ** *P* < 0.01; ****, *P* < 0.0001; Mann-Whitney test). (**F**) Analysis of chromatin binding of FANCJ in MRC5SV and shWRNIP1 cells. Chromatin fraction of cells, treated or not with the proteasome inhibitor MG132 for 4 h, were analysed by immunoblotting. The membrane was blotted with anti-FANCJ antibody. LAMIN B1 was used as a loading for the chromatin fraction. The amount of the chromatin-bound FANCJ is reported as ratio of FANCJ/LAMIN B1 normalized over the untreated control. (**G**) Analysis of chromatin binding of FANCJ in MRC5SV and shWRNIP1 cells. Chromatin fraction of cells treated or not with 0.4 µM Aph for 24 h and the transcription inhibitor DRB the last three hours were analysed by immunoblotting as in (D).

Next, we wondered if G4s localised on parental ssDNA upon Aph-induced MRS. Using a fluorescence-based PLA assay, we investigated the co-localisation of G4s at/near ssDNA. To do this, cells were labelled with IdU as in Figure 7A, then subjected to PLA assay. As showed in Figure 7B, we found that the co-localisation between ssDNA (anti-IdU signal) and G4s (anti-BG4 signal) significantly increased after Aph in both cell lines tested, but the number of spots was higher in the absence of WRNIP1 than in wild-type counterparts (Figure 7B). Moreover, DRB greatly suppressed the G4-ssDNA interaction, suggesting a strong association between G4s and R-loops in shWRNIP1 cells upon MRS.

### WRNIP1 and FANCJ interacts in response to MRS

Recently, Mgs1, the yeast homologue of WRNIP1, has been identified as a novel G4-binding protein that, supporting replication at G4-forming genomic regions, participates in the maintenance of genome integrity ^34^. Hence, we wondered whether WRNIP1 resided at/near G4s upon MRS. Consistently with data in yeast ^34^, our PLA assay revealed a co-localization of WRNIP1 with G4-structures at comparable levels in shWRNIP1^WT^ and shWRNIP1^T294A^ cells under unperturbed and Aph-treated conditions but higher in shWRNIP1^D37A^, confirming that, also in human cells, loss of ATPase domain of WRNIP1 does not affect this interaction (Figure 7C).

In human cells, FANCJ is the most potent G4-structure-unwinding helicases ^23^, thus, we investigated if WRNIP1 and FANCJ interact *in vivo* testing their co-immunoprecipitation. To this aim, HEK293T cells were transfected with an empty vector or the FLAG-tagged wild-type WRNIP1 and treated or not with Aph or PDS. Western blot showed that WRNIP1 and FANCJ immunoprecipitated under untreated conditions, and that Aph, more than PDS, slightly increased this interaction (Figure 7D). This result supports a possible cooperation of WRNIP1 and FANCJ in response to Aph-induced MRS. PLA assay corroborated this interaction and revealed that increased number of spots induced by Aph in shWRNIP1^WT^ cells was reduced by transcription inhibition (Figure 7E). Notably, loss of UBZ domain of WRNIP1 compromised WRNIP1 and FANCJ interaction (Figure 7E).

Next, we measured the levels of FANCJ in WRNIP1-deficient cells. Western blot analysis after cellular fractionation showed that, under unperturbed conditions, the amount of chromatin-bound FANCJ was lower in shWRNIP1 than in wild-type cells (Figure 7F). Notably, MG132 treatment demonstrated that FANCJ was susceptible to proteasome-mediated degradation, suggesting the possibility that FANCJ might not be properly stabilized in the absence of WRNIP1. While, in wild-type cells, Aph treatment promoted chromatin loading of FANCJ that was modulated by DRB, in WRNIP1-deficient cells led to FANCJ halved levels (Figure 7G). Interestingly, chromatin loading of FANCJ seemed affected by loss of UBZ domain of WRNIP1 but not of its ATPase activity (Suppl. Figure 8A). Similar results were obtained treating cells with PDS (Suppl. Figure 8B). Furthermore, defective FANCJ stabilization in WRNIP1-deficient cells was independent of cell type (Suppl. Figure 8C).

These results suggest a cooperation between WRNIP1 and FANCJ in the response to MRS.

### FANCJ is needed for counteracting R-loop-dependent G4 formation in cells lacking WRNIP1 or its UBZ domain upon MRS

Previous study revealed that G4-stabilizing ligand increases the requirement for the DNA helicase FANCJ ^53^. Accordingly, depletion of FANCJ causes G4 accumulation, DNA damage and persistent replication stalling at G4s ^54^, demonstrating a critical role for the helicase in resolving these structures. Hence, we explored the consequence of concomitant loss of WRNIP1 and FANCJ for G4 accumulation and DNA damage upon MRS. To this aim, we evaluated the levels of G4s in MRC5SV and shWRNIP1 cells depleted of FANCJ by RNAi, treated or not with Aph (Figure 8A). Immunofluorescence analysis confirmed that loss of WRNIP1 or FANCJ resulted in a pronounced spontaneous increase in BG4 nuclear intensity that was significantly enriched by treatment (Figure 8A). Interestingly, concomitant loss of WRNIP1 and FANCJ did not exacerbate the phenotype exhibited by the single deficiencies (Figure 8A). In agreement with this, loss of FANCJ in wild-type or shWRNIP1 cells resulted in a similar amount of Aph-induced DNA damage (Suppl. Figure 9A and B). Furthermore, PLA assay demonstrated an Aph- and transcription-dependent co-localization between FANCJ and G4s, which was reduced in the absence of WRNIP1 but restored via stabilising FANCJ by MG132 treatment (Figure 8B and Suppl. Figure 10). By contrast, WRNIP1/G4s co-localization was not affected by loss of FANCJ (Figure 8C). Notably, stabilisation of FANCJ led to a significant lowering of the pronounced BG4 nuclear intensity detected in WRNIP1-deficient cells upon Aph (Figure 8D).

**Figure 8.**
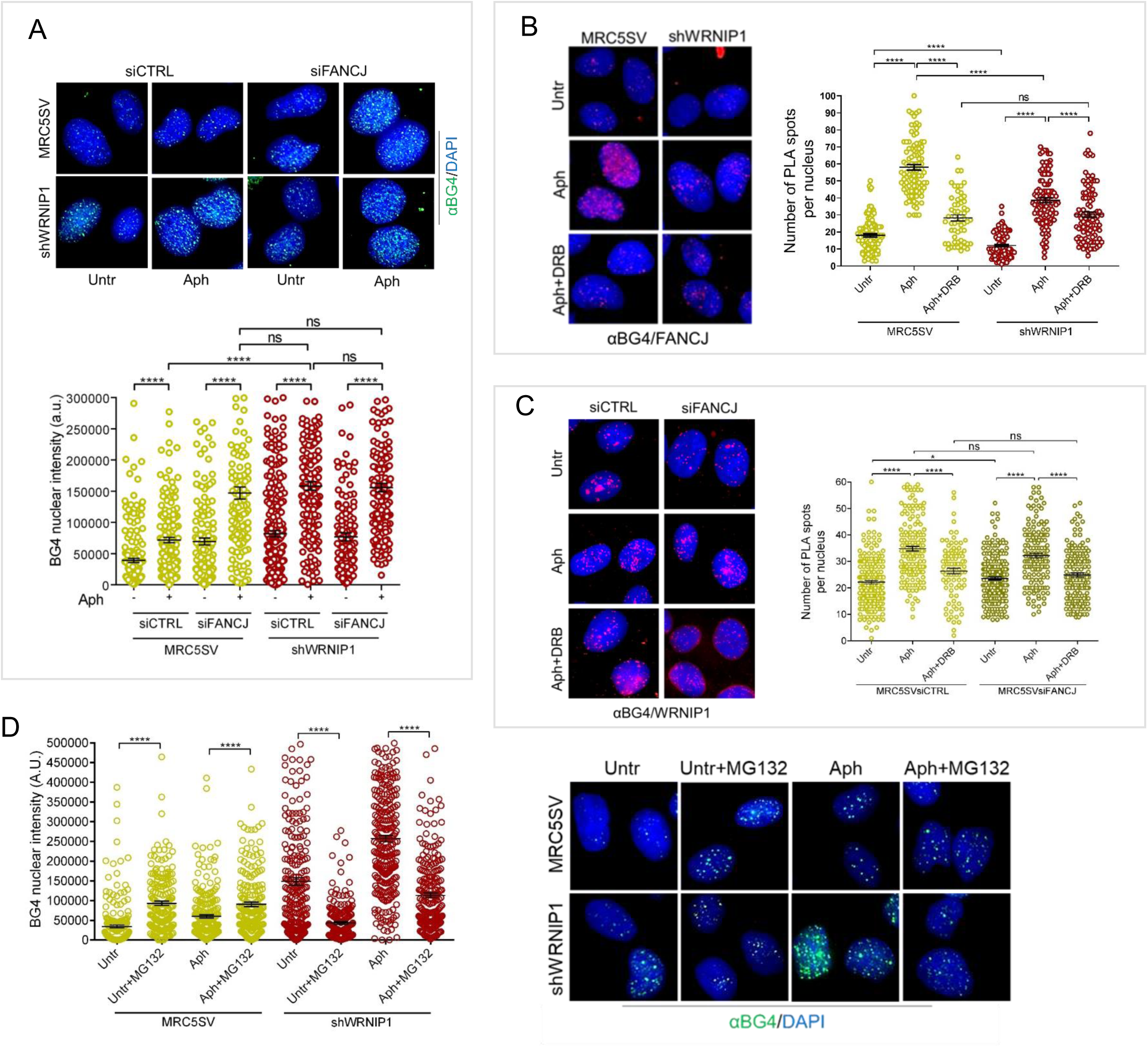
**Evaluation of the interplay between WRNIP1 and FANCJ in G4 resolution** (**A**) Effect of FANCJ deregulation on G4s levels in MRC5SV and shWRNIP1cells treated or not with Aph 0.4 µM Aph for 24 h. Representative images are given. Nuclei were counterstained with DAPI. The graph shows nuclear BG4 fluorescence intensity. Data are presented as means of three independent experiments. Horizontal black lines represent the mean. Error bars represent standard error (ns, not significant; ****, *P* < 0.0001; Mann-Whitney test). (**B**) Evaluation of FANCJ and G4 co-localization by PLA assay in MRC5SV and shWRNIP1 cells treated or not with 0.4 µM Aph for 24 h and the transcription inhibitor DRB the last 3 h. Representative images are given. Each red spot represents a single interaction between G4s and FANCJ. Nuclei were counterstained with DAPI. Dot plot shows the number of PLA spots per nucleus. Data are presented as means of three independent experiments. Horizontal black lines represent the mean ± SE (ns, not significant; ****, *P* < 0.0001; Mann-Whitney test). (**C**) PLA assay to evaluate WRNIP1 recruitment to G4s in the absence of FANCJ in MRC5SV cells treated or not with 0.4 µM Aph for 24 h alone or in combination with DRB 50 µM for the last 3 h. Cells were fixed and stained with antibodies against WRNIP1 and G4s. Representative images are given. Each red spot represents a single interaction between protein and DNA. Nuclei were counterstained with DAPI. Dot plot shows the number of PLA spots per nucleus. Data are presented as means of three independent experiments. Horizontal black lines represent the mean ± SE (ns, not significant; * *P* < 0.05; ****, *P* < 0.0001; Mann-Whitney test). (**D**) Immunostaining with BG4 in shWRNIP1^WT^and shWRNIP1 cells treated or not with 0.4 µM Aph for 24 h in combination with the proteasome inhibitor MG132 for the last 6 h. Visualization of G4s was carried out as previously described. Representative images are given. The graph shows the quantification of BG4 nuclear intensity per nucleus. Data are presented as means of three independent experiments. Horizontal black lines represent the mean ± SE (****, *P* < 0.0001; Mann-Whitney test).

Overall, these findings suggest that WRNIP1 may contribute to stabilisation of FANCJ to G4 sites upon MRS.

## DISCUSSION

Discovered as one of the interactors of the WRN helicase ^55, 58^, the precise function of WRNIP1 in human cells is largely unknown. Previous results from our lab and others, showed that WRNIP1 is crucial for the protection of HU-stalled replication forks ^28, 29^. In this study, we uncover a role for WRNIP1 and its ubiquitin-binding (UBZ) domain in counteracting transcription-replication conflicts (TRCs) associated with persistent G4/R-loops when replication is perturbed. Furthermore, we identify an interplay between WRNIP1 and the DNA helicase FANCJ in dealing with G4 formation at R-loops after mild replication stress (MRS).

More recently, it has been shown that checkpoint defects elicit a WRNIP1-mediated response to limit R-loop-associated genomic instability ^31^. Although R-loops, the main transcription-associated DNA secondary structures, regulate several physiological processes, they may be potent replication fork barriers ^1, 3^, and accumulation of aberrant R-loops leads to DNA damage in both yeast and human cells ^4, 61, 64, 67, 70^. Here, we demonstrate that loss of WRNIP1 causes transcription-dependent DNA damage and impairs replication fork progression not only after perturbed replication but already under unperturbed conditions. Of note, we observe that, although the ATPase activity of WRNIP1 is important for the restart of stalled forks ^28^, it is dispensable to cope with TRCs. Aside from its ATPase activity, WRNIP1 possesses an UBZ domain with undefined function after replication perturbation^25^. In contrast to loss of the ATPase activity, a mutation that disrupts the WRNIP1 UBZ domain ^35, 36^ is sufficient to stimulate TRCs and DNA damage at levels that are comparable if not higher to what observed in WRNIP1-deficient cells.

Persistence of R-loops is deleterious because they may interfere with DNA replication, and R-loop-mediated fork stalling is a major feature of TRCs ^39, 62, 65, 68^. In WRNIP1-deficient or WRNIP1 UBZ mutant cells, the elevated DNA damage and reduced fork speed are reverted by RNase H1-GFP overexpression, which is known to degrade RNA/DNA hybrids and wipes out R-loops ^44^. In line with this, much more R-loops are detected in WRNIP1-deficient or WRNIP1 UBZ mutant cells. This suggests that persistent R-loops are likely responsible for the increased levels of TRCs, the marked replication defects and genomic instability observed in the absence of WRNIP1.

R-loop is a three-stranded nucleic acid structure that consists of a DNA/RNA hybrid and a displaced strand of DNA. It is noteworthy that the displaced ssDNA of R-loop has the propensity to fold into G4s, which stabilize the R-loop and promote its extension ^19, 20^. Our analyses demonstrate that loss of WRNIP1 or its functional UBZ domain causes an R-loop-dependent G4 formation. While the precise mechanism of their formation, in our context, remains to be defined, our data show that G4s and R-loops co-localize in a transcription-dependent manner upon MRS. Moreover, our immunofluorescence analysis places G4s at/near ssDNA of R-loop. Together these findings support the hypothesis that G4s detected in WRNIP1-deficient and WRNIP1 UBZ mutant cells reside mainly within R-loops.

The yeast homolog of WRNIP1, Mgs1 exhibits a high G4-binding affinity without requiring its ATPase activity ^34^. Consistently with data in yeast, we reveal a co-localization of WRNIP1 to G4s upon MRS that does not require ATPase activity but is improved by loss of WRNIP1 UBZ domain. Although G4s have a positive function in the genome, their misregulation can affect different cellular processes ^71, 73, 74^. The current model proposes that G4s, which form during DNA replication, slow down moving forks ^75^. Consistently, our findings suggest that G4s accumulation in the absence of WRNIP1 or a functional UBZ domain depends on replication, and that replication defects and genomic instability are primarily caused by TRCs deriving from inefficient removal of G4/R-loops. In agreement with this, we find that a G4-stabilizing ligand enhances TRCs in WRNIP1-deficient or WRNIP1 UBZ mutant cells that can be alleviated by pre-treatment with inhibitors of both the replication and transcription.

How does WRNIP1 contribute to prevent G4 accumulation at R-loops and thus promote fork progression? WRNIP1 promotes stalled fork restart ^28^. However, WRNIP1-dependent stimulation of fork restart apparently requires its ATPase function ^28^, while protection from TRCs associated with G4/R-loops does not. In contrast, the UBZ domain of WRNIP1 is critical for the prevention of this kind of TRCs. As R-loop homeostasis is regulated by ubiquitination ^56^, and given that WRNIP1 can interact with different ubiquitin signals ^27^, one possible explanation is that WRNIP1 may play a role in an ubiquitination-dependent mechanism for the resolution of G4/R-loop. WRNIP1 and its UBZ domain have been implicated previously as “recruiting docks” for proteins during the checkpoint response after DNA damage ^59^ and ubiquitin-binding is often exploited by proteins acting at fork or during DNA repair ^63^. Thus, WRNIP1 might contribute to the recruitment of other factors involved in G4s resolution during the DNA replication.

It is known that G4 structures are unfolded by helicases ^22, 23^. In yeast, Pif1 helicase stimulates Mgs1 recruitment to G4s ^34^. Although how Mgs1 and Pif1 interact and work together are not completely clarified, it has been hypothesized that they may cooperate in G4 unwinding or by promoting replication restart or controlling/stabilizing the replication fork at G4 sites ^34^. In human cells, several G4-resolving DNA helicases exists, including the 5′→3′ helicase FANCJ ^23, 53, 66^ or the WRN and BLM proteins, with opposing directionality ^69^. However, FANCJ is considered the most effective unwinder of G4 structures in human cells ^23^. Similar to WRNIP1, FANCJ suppresses accumulation of G4 structures within replisomes in human cells ^54^, and a role in promoting DNA synthesis past G4s has been demonstrated in Xenopus egg extracts ^53^. In agreement with a role of FANCJ in DNA replication, its deficiency leads to G4 accumulation, DNA damage at G4-associated replication forks upon MRS ^54^, all phenotypes shared with WRNIP1-deficient or UBZ-mutated cells. However, it is not clear how FANCJ is recruited to the sites of G4s at stalled forks.

Unlike in yeast, we see that WRNIP1 and FANCJ physically interact under either unperturbed or stressful conditions. Notably, WRNIP1-deficient cells exhibit defective interaction between FANCJ and G4s. By contrast, WRNIP1 and G4s co-localization is a FANCJ-independent event. Hence, we propose that WRNIP1 is necessary to recruit FANCJ or maintain it stable at sites of G4s. Indeed, the level of chromatin-bound FANCJ is low in the absence of WRNIP1 or its UBZ domain. FANCJ levels can be rescued by the proteasome inhibitor MG132 in the absence of WRNIP1, suggesting that WRNIP1 once associated with FANCJ stabilises it in chromatin. Interestingly, since WRNIP1-deficient cells regain the ability to reduce G4s by counteracting FANCJ degradation, it is very likely that WRNIP1 and its UBZ domain-dependent stabilization of the DNA helicase represents a novel mechanism to promote G4s resolution. From this point of view, it is plausible that defective fork progression is directly correlated to inability to support FANCJ in G4 resolution. WRNIP1 also interacts with DNA polymerase delta ^72^, suggesting that this under-investigated protein might be a co-factor for multiple replication-related proteins under replication stress.

In summary, our findings uncover a novel mechanism in counteracting pathological accumulation of G4/R-loops, leading to increased genomic instability upon MRS in human cells. As mounting evidence reveals that G4/R-loops may mediated genomic instability observed in human cancer cells ^19^, and given that WRNIP1 was found overexpressed in the most common cancers worldwide, such as lung and breast cancers, understanding the mechanisms required for the prevention of TRCs could, therefore, open new avenues in the definition of different pathways leading to genomic instability in cancer.

## MATERIALS AND METHODS

### Cell lines and culture conditions

The SV40-transformed MRC5 fibroblast cell line (MRC5SV) was a generous gift from Patricia Kannouche (IGR, Villejuif, France). MRC5SV cells stably expressing WRNIP1-targeting shRNA (shWRNIP1) and isogenic cell lines stably expressing the RNAi-resistant full-length wild-type WRNIP1 (shWRNIP1^WT^), its ATPase-dead mutant form (shWRNIP1^T294A^) were generated as previous reported ^28^. By using the Neon^TM^ Transfection System Kit (Invitrogen) according to the manufacturer’s instructions, shWRNIP1 cells were stably transfected with a plasmid expressing a FLAG-tagged full-length WRNIP1 plasmid carrying Ala substitution at Asp37 site missense-mutant form of WRNIP1 with dead form of ubiquitin-binding zinc finger (UBZ) domain (WRNIP1^D37A^) ^25^, Cells were cultured in the presence of neomycin and puromycin (1 mg/ml and 100 ng/ml, respectively) to maintain selective pressure for expression.

The human embryonic kidney 293 cells (HEK293T) and the human osteosarcoma U2OS cells were obtained from American Type Culture Collection (VA, USA).

All cell lines were maintained in DMEM media supplemented with 10% FBS, 100 U/mL penicillin and 100 mg/mL streptomycin and incubated at 37°C in an humified 5% CO2 atmosphere.

### Immunofluorescence

Immunofluorescence analysis for γ-H2AX was performed as previously described ^57^. Briefly, exponential growing cells were seeded onto Petri dish, then treated (or mock-treated) as indicated, fixed in % formaldehyde for 10 min, and permeabilized using 0.4% Triton X-100 for 10 min before being incubated with 10% FBS for 1 h at RT. After blocking, for γ-H2AX detection, cells were incubated with the following primary antibody: mouse-monoclonal anti-γ-H2AX (Millipore, 1:1000). Immunofluorescence for G-quadruplex structures (G4s) was performed as described ^60^ with minor changes. Cells grown on glass coverslips were fixed with ice-cold 80% methanol in PBS for 15 min at -20°C, then washed two times in PBS. Next, cells were blocked with 10% FBS/PBS for 1 h and incubated with the anti-G-quadruplex antibody (BG4, Sigma-Aldrich, 1:200) overnight at 4°C.

Immunostaining for RNA-DNA hybrids was performed as described ^17^ . Briefly, cells were fixed in 100% methanol for 10 min at -20°C, washed three times in PBS, pre-treated with 6 μg/ml of RNase A for 45 min at 37°C in 10 mM Tris-HCl pH 7.5 supplemented with 0.5 M NaCl, before blocking in 2% BSA/PBS overnight at 4°C. Cells were then incubated with the anti-DNA-RNA hybrid [S9.6] antibody (Kerafast, 1:100) overnight at 4°C.

To detect parental-strand ssDNA, cells were pre-labelled for 20 h with 100 µM IdU (Sigma-Aldrich), washed in drug-free medium for 2 h and then treated with Aph for 24 h. Next, cells were washed with PBS, permeabilized with 0.5% Triton X-100 for 10 min at 4°C, fixed with 3% formaldehyde/2% sucrose solution for 10 min, and then blocked in 3% BSA/PBS for 15 min as previously described ^28^. Fixed cells were then incubated with anti-IdU antibody (mouse-monoclonal antiBrdU/IdU; clone b44 Becton Dickinson, 1:100).

After each primary antibody, cells were washed twice with PBS, and incubated with the specific secondary antibody: goat anti-mouse Alexa Fluor-488 or goat anti-rabbit Alexa Fluor-594 (Molecular Probes). The incubation with secondary antibodies were accomplished in a humidified chamber for 1 h at RT. DNA was counterstained with 0.5 μg/ml DAPI.

Images were acquired randomly using Eclipse 80i Nikon Fluorescence Microscope, equipped with a VideoConfocal (ViCo) system. For each time point, at least 200 nuclei were examined. Nuclear foci were scored at a 40× magnification and only nuclei showing more than five bright foci were counted as positive. Intensity per nucleus was calculated using ImageJ. Parallel samples incubated with either the appropriate normal serum or only with the secondary antibody confirmed that the observed fluorescence pattern was not attributable to artefacts.

### Dot blot analysis

Dot blot analysis was performed according to the protocol reported elsewhere ^45^. Genomic DNA was isolated by standard extraction with Phenol/Clorophorm/Isoamylic Alcohol (pH 8.0) followed by precipitation with 3 M NaOAc and 70% Ethanol. Isolated gDNA was randomly fragmented overnight at 37°C with a cocktail of restriction enzymes (BsrgI, EcoRI, HindIII, XbaI) supplemented with 1 M Spermidin. After incubation, digested DNA was cleaned up with Phenol/Clorophorm extraction and standard Ethanol precipitation. After sample quantification, 5 μg of digested DNA were incubated with RNaseH overnight at 37°C as a negative control. Five micrograms of each sample were spotted onto a nitrocellulose membrane, blocked in 5% non-fat dry milk and incubated with the anti-DNA-RNA hybrid [S9.6] antibody (Kerafast, 1:1000) overnight at 4°C. Horseradish peroxidase-conjugated goat specie-specific secondary antibody (Santa Cruz Biotechnology, Inc.) was used. Quantification on scanned image of blot was performed using Image Lab software.

### Statistical analysis

Statistical analysis was performed using Prism 8 (GraphPad Software). Details of the individual statistical tests are indicated in the figure legends and results. Statistical differences in all case were determined by Student’s t-test or Mann-Whitney test. In all cases, not significant; *P* > 0.05; * *P* < 0.05; ** *P* < 0.01; *** *P* < 0.001; **** *P* < 0.0001. All experiments were repeated at least three times unless otherwise noted.

## FUNDING

This research was funded by Associazione Italiana per la Ricerca sul Cancro to A.F. (IG #19971) and to P.P. (IG #17383).

## Notes

### Competing Interest Statement

The authors have declared no competing interest.

## REFERENCES

1. Gómez-González, B., and Aguilera, A. (2019). Transcription-mediated replication hindrance: A major driver of genome instability. Genes Dev. 33, 1008–1026. 10.1101/gad.324517.119.

2. García-Muse, T., and Aguilera, A. (2019). R Loops: From Physiological to Pathological Roles. Cell 179, 604–618. 10.1016/j.cell.2019.08.055.

3. Crossley, M.P., Bocek, M., and Cimprich, K.A. (2019). R-Loops as Cellular Regulators and Genomic Threats. Mol. Cell 73, 398–411. 10.1016/j.molcel.2019.01.024.

4. Santos-Pereira, J.M., and Aguilera, A. (2015). R loops: New modulators of genome dynamics and function. Nat. Rev. Genet. 16, 583–597. 10.1038/nrg3961.

5. Brambati, A., Zardoni, L., Nardini, E., Pellicioli, A., and Liberi, G. (2020). The dark side of RNA:DNA hybrids. Mutat. Res. - Rev. Mutat. Res. 784, 108300. 10.1016/j.mrrev.2020.108300.

6. Sanz, L.A., Hartono, S.R., Lim, Y.W., Steyaert, S., Rajpurkar, A., Ginno, P.A., Xu, X., and Chédin, F. (2016). Prevalent, Dynamic, and Conserved R-Loop Structures Associate with Specific Epigenomic Signatures in Mammals. Mol. Cell 63, 167–178. 10.1016/j.molcel.2016.05.032.

7. Wahba, L., Amon, J.D., Koshland, D., and Vuica-Ross, M. (2011). RNase H and Multiple RNA Biogenesis Factors Cooperate to Prevent RNA:DNA Hybrids from Generating Genome Instability. Mol. Cell 44, 978–988. 10.1016/j.molcel.2011.10.017.

8. Helmrich, A., Ballarino, M., and Tora, L. (2011). Collisions between Replication and Transcription Complexes Cause Common Fragile Site Instability at the Longest Human Genes. Mol. Cell 44, 966–977. 10.1016/j.molcel.2011.10.013.

9. Shivji, M.K.K., Renaudin, X., Williams, Ç.H., and Venkitaraman, A.R. (2018). BRCA2 Regulates Transcription Elongation by RNA Polymerase II to Prevent R-Loop Accumulation. Cell Rep. 22, 1031–1039. 10.1016/j.celrep.2017.12.086.

10. Hatchi, E., Skourti-Stathaki, K., Ventz, S., Pinello, L., Yen, A., Kamieniarz-Gdula, K., Dimitrov, S., Pathania, S., McKinney, K.M., Eaton, M.L., et al. (2015). BRCA1 recruitment to transcriptional pause sites is required for R-loop-driven DNA damage repair. Mol. Cell 57, 636–647. 10.1016/j.molcel.2015.01.011.

11. Schwab, R.A., Nieminuszczy, J., Shah, F., Langton, J., Lopez Martinez, D., Liang, C.C., Cohn, M.A., Gibbons, R.J., Deans, A.J., and Niedzwiedz, W. (2015). The Fanconi Anemia Pathway Maintains Genome Stability by Coordinating Replication and Transcription. Mol. Cell 60, 351–361. 10.1016/j.molcel.2015.09.012.

12. García-Rubio, M.L., Pérez-Calero, C., Barroso, S.I., Tumini, E., Herrera-Moyano, E., Rosado, I. V., and Aguilera, A. (2015). The Fanconi Anemia Pathway Protects Genome Integrity from R-loops. PLoS Genet. 11, 1–17. 10.1371/journal.pgen.1005674.

13. Bhatia, V., Barroso, S.I., García-Rubio, M.L., Tumini, E., Herrera-Moyano, E., and Aguilera, A. (2014). BRCA2 prevents R-loop accumulation and associates with TREX-2 mRNA export factor PCID2. Nature 511, 362–365. 10.1038/nature13374.

14. Liang, Z., Liang, F., Teng, Y., Chen, X., Liu, J., Longerich, S., Rao, T., Green, A.M., Collins, N.B., Xiong, Y., et al. (2019). Binding of FANCI-FANCD2 Complex to RNA and R-Loops Stimulates Robust FANCD2 Monoubiquitination. Cell Rep. 26, 564–572.e5. 10.1016/j.celrep.2018.12.084.

15. Urban, V., Dobrovolna, J., Hühn, D., Fryzelkova, J., Bartek, J., and Janscak, P. (2016). RECQ5 helicase promotes resolution of conflicts between replication and transcription in human cells. J. Cell Biol. 214, 401–415. 10.1083/jcb.201507099.

16. Chang, E.Y.C., Novoa, C.A., Aristizabal, M.J., Coulombe, Y., Segovia, R., Chaturvedi, R., Shen, Y., Keong, C., Tam, A.S., Jones, S.J.M., et al. (2017). RECQ-like helicases Sgs1 and BLM regulate R-loop-associated genome instabil. J. Cell Biol. 216, 3991–4005. 10.1083/jcb.201703168.

17. Marabitti, V., Lillo, G., Malacaria, E., Palermo, V., Sanchez, M., Pichierri, P., and Franchitto, A. (2019). ATM pathway activation limits R-loop-associated genomic instability in Werner syndrome cells. Nucleic Acids Res. 47, 3485–3502. 10.1093/nar/gkz025.

18. Tresini, M., Warmerdam, D.O., Kolovos, P., Snijder, L., Vrouwe, M.G., Demmers, J.A.A., Van Ijcken, W.F.J., Grosveld, F.G., Medema, R.H., Hoeijmakers, J.H.J., et al. (2015). The core spliceosome as target and effector of non-canonical ATM signalling. Nature 523, 53–58. 10.1038/nature14512.

19. De Magis, A., Manzo, S.G., Russo, M., Marinello, J., Morigi, R., Sordet, O., and Capranico, G. (2019). DNA damage and genome instability by G-quadruplex ligands are mediated by R loops in human cancer cells. Proc. Natl. Acad. Sci. U. S. A. 116, 816–825. 10.1073/pnas.1810409116.

20. Duquette, M.L., Handa, P., Vincent, J.A., Taylor, A.F., and Maizels, N. (2004). Intracellular transcription of G-rich DNAs induces formation of G-loops, novel structures containing G4 DNA. Genes Dev. 18, 1618–1629. 10.1101/gad.1200804.

21. Spiegel, J., Adhikari, S., and Balasubramanian, S. (2020). The Structure and Function of DNA G-Quadruplexes. Trends Chem. 2, 123–136. 10.1016/j.trechm.2019.07.002.

22. Mendoza, O., Bourdoncle, A., Boulé, J.B., Brosh, R.M., and Mergny, J.L. (2016). G-quadruplexes and helicases. Nucleic Acids Res. 44, 1989–2006. 10.1093/nar/gkw079.

23. Sauer, M., and Paeschke, K. (2017). G-quadruplex unwinding helicases and their function in vivo. Biochem. Soc. Trans. 45, 1173–1182. 10.1042/BST20170097.

24. Brosh, R.M., and Wu, Y. (2021). An emerging picture of FANCJ’s role in G4 resolution to facilitate DNA replication. NAR Cancer 3, 1–8. 10.1093/narcan/zcab034.

25. Yoshimura, A., Seki, M., and Enomoto, T. (2017). The role of WRNIP1 in genome maintenance. Cell Cycle 16, 515–521. 10.1080/15384101.2017.1282585.

26. Yoshimura, A., Seki, M., Kanamori, M., Tateishi, S., Tsurimoto, T., Tada, S., and Enomoto, T. (2009). Physical and functional interaction between WRNIP1 and RAD18. Genes Genet. Syst. 84, 171–178. 10.1266/ggs.84.171.

27. Crosetto, N., Bienko, M., Hibbert, R.G., Perica, T., Ambrogio, C., Kensche, T., Hofmann, K., Sixma, T.K., and Dikic, I. (2008). Human Wrnip1 is localized in replication factories in a ubiquitin-binding zinc finger-dependent manner. J. Biol. Chem. 283, 35173–35185. 10.1074/jbc.M803219200.

28. Leuzzi, G., Marabitti, V., Pichierri, P., and Franchitto, A. (2016). WRNIP 1 protects stalled forks from degradation and promotes fork restart after replication stress . EMBO J. 35, 1437–1451. 10.15252/embj.201593265.

29. Porebski, B., Wild, S., Kummer, S., Scaglione, S., Gaillard, P.H.L., and Gari, K. (2019). WRNIP1 Protects Reversed DNA Replication Forks from SLX4-Dependent Nucleolytic Cleavage. iScience 21, 31–41. 10.1016/j.isci.2019.10.010.

30. Kanu, N., Zhang, T., Burrell, R.A., Chakraborty, A., Cronshaw, J., Dacosta, C., Grönroos, E., Pemberton, H.N., Anderton, E., Gonzalez, L., et al. (2016). RAD18, WRNIP1 and ATMIN promote ATM signalling in response to replication stress. Oncogene 35, 4009–4019. 10.1038/onc.2015.427.

31. Marabitti, V., Lillo, G., Malacaria, E., Palermo, V., Pichierri, P., and Franchitto, A. (2020). Checkpoint defects elicit a WRNIP1-mediated response to counteract R-loop-associated genomic instability. Cancers (Basel). 12. 10.3390/cancers12020389.

32. Pladevall-Morera, D., Munk, S., Ingham, A., Garribba, L., Albers, E., Liu, Y., Olsen, J. V., and Lopez-Contreras, A.J. (2019). Proteomic characterization of chromosomal common fragile site (CFS)-associated proteins uncovers ATRX as a regulator of CFS stability. Nucleic Acids Res. 47, 8004–8018. 10.1093/nar/gkz510.

33. Hishida, T., Iwasaki, H., Ohno, T., Morishita, T., and Shinagawa, H. (2001). A yeast gene, MGS1, encoding a DNA-dependent AAA+ ATPase is required to maintain genome stability. Proc. Natl. Acad. Sci. U. S. A. 98, 8283–8289. 10.1073/pnas.121009098.

34. Zacheja, T., Toth, A., Harami, G.M., Yang, Q., Schwindt, E., Kovács, M., Paeschke, K., and Burkovics, P. (2020). Mgs1 protein supports genome stability via recognition of G-quadruplex DNA structures. FASEB J. 34, 12646–12662. 10.1096/fj.202000886R.

35. Bish, R.A., and Myers, M.P. (2007). Werner helicase-interacting protein 1 binds polyubiquitin via its zinc finger domain. J. Biol. Chem. 282, 23184–23193. 10.1074/jbc.M701042200.

36. Nomura, H., Yoshimura, A., Edo, T., Kanno, S.I., Tada, S., Seki, M., Yasui, A., and Enomoto, T. (2012). WRNIP1 accumulates at laser light irradiated sites rapidly via its ubiquitin-binding zinc finger domain and independently from its ATPase domain. Biochem. Biophys. Res. Commun. 417, 1145–1150. 10.1016/j.bbrc.2011.12.080.

37. Müller, W.E., Seibert, G., Beyer, R., Breter, H.J., Maidhof, A., and Zahn, R.K. (1977). Effect of Cordycepin on Nucleic Acid Metabolism in L5178Y Cells and on Nucleic Acid-synthesizing Enzyme Systems. Cancer Res. 37, 3824–3833.

38. Ward, I.M., and Chen, J. (2001). Histone H2AX Is Phosphorylated in an ATR-dependent Manner in Response to Replicational Stress. J. Biol. Chem. 276, 47759–47762. 10.1074/jbc.C100569200.

39. Gan, W., Guan, Z., Liu, J., Gui, T., Shen, K., Manley, J.L., and Li, X. (2011). R-loop-mediated genomic instability is caused by impairment of replication fork progression. Genes Dev. 25, 2041–2056. 10.1101/gad.17010011.

40. Allison, D.F., and Wang, G.G. (2019). R-loops: Formation, function, and relevance to cell stress. Cell Stress 3, 38–46. 10.15698/cst2019.02.175.

41. Gaillard, H., and Aguilera, A. (2016). Transcription as a Threat to Genome Integrity. Annu. Rev. Biochem. 85, 291–317. 10.1146/annurev-biochem-060815-014908.

42. Boguslawski, S.J., Smith, D.E., Michalak, M.A., Mickelson, K.E., Yehle, C.O., Patterson, W.L., and Carrico, R.J. (1986). Characterization of monoclonal antibody to DNA · RNA and its application to immunodetection of hybrids. J. Immunol. Methods 89, 123–130. 10.1016/0022-1759(86)90040-2.

43. Hamperl, S., Bocek, M.J., Saldivar, J.C., Swigut, T., and Cimprich, K.A. (2017). Transcription-Replication Conflict Orientation Modulates R-Loop Levels and Activates Distinct DNA Damage Responses. Cell 170, 774–786.e19. 10.1016/j.cell.2017.07.043.

44. Cerritelli, S.M., Frolova, E.G., Feng, C., Grinberg, A., Love, P.E., and Crouch, R.J. (2003). Failure to produce mitochondrial DNA results in embryonic lethality in Rnaseh1 null mice. Mol. Cell 11, 807–815. 10.1016/S1097-2765(03)00088-1.

45. Morales, J.C., Richard, P., Patidar, P.L., Motea, E.A., Dang, T.T., Manley, J.L., and Boothman, D.A. (2016). XRN2 Links Transcription Termination to DNA Damage and Replication Stress. PLoS Genet. 12, 1–22. 10.1371/journal.pgen.1006107.

46. García-Muse, T., and Aguilera, A. (2016). Transcription-replication conflicts: How they occur and how they are resolved. Nat. Rev. Mol. Cell Biol. 17, 553–563. 10.1038/nrm.2016.88.

47. Söderberg, O., Leuchowius, K.J., Gullberg, M., Jarvius, M., Weibrecht, I., Larsson, L.G., and Landegren, U. (2008). Characterizing proteins and their interactions in cells and tissues using the in situ proximity ligation assay. Methods 45, 227–232. 10.1016/j.ymeth.2008.06.014.

48. Belotserkovskii, B.P., Tornaletti, S., D’Souza, A.D., and Hanawalt, P.C. (2018). R-loop generation during transcription: Formation, processing and cellular outcomes. DNA Repair (Amst). 71, 69–81. 10.1016/j.dnarep.2018.08.009.

49. Biffi, G., Tannahill, D., McCafferty, J., and Balasubramanian, S. (2013). Quantitative visualization of DNA G-quadruplex structures in human cells. Nat. Chem. 5, 182–186. 10.1038/nchem.1548.

50. Koirala, D., Dhakal, S., Ashbridge, B., Sannohe, Y., Rodriguez, R., Sugiyama, H., Balasubramanian, S., and Mao, H. (2011). A single-molecule platform for investigation of interactions between G-quadruplexes and small-molecule ligands. Nat. Chem. 3, 782–787. 10.1038/nchem.1126.

51. Rodriguez, R., Miller, K.M., Forment, J. V., Bradshaw, C.R., Nikan, M., Britton, S., Oelschlaegel, T., Xhemalce, B., Balasubramanian, S., and Jackson, S.P. (2012). Small-molecule-induced DNA damage identifies alternative DNA structures in human genes. Nat. Chem. Biol. 8, 301– 310. 10.1038/nchembio.780.

52. Basile, G., Leuzzi, G., Pichierri, P., and Franchitto, A. (2014). Checkpoint-dependent and independent roles of the Werner syndrome protein in preserving genome integrity in response to mild replication stress. Nucleic Acids Res. 42, 12628–12639. 10.1093/nar/gku1022.

53. Castillo Bosch, P., Segura-Bayona, S., Koole, W., Heteren, J.T., Dewar, J.M., Tijsterman, M., and Knipscheer, P. (2014). FANCJ promotes DNA synthesis through G-quadruplex structures . EMBO J. 33, 2521–2533. 10.15252/embj.201488663.

54. Lee, W.T.C., Yin, Y., Morten, M.J., Tonzi, P., Gwo, P.P., Odermatt, D.C., Modesti, M., Cantor, S.B., Gari, K., Huang, T.T., et al. (2021). Single-molecule imaging reveals replication fork coupled formation of G-quadruplex structures hinders local replication stress signaling. Nat. Commun. 12. 10.1038/s41467-021-22830-9.

55. Kawabe, Y.I., Branzei, D., Hayashi, T., Suzuki, H., Masuko, T., Onoda, F., Heo, S.J., Ikeda, H., Shimamoto, A., Furuichi, Y., et al. (2001). A Novel Protein Interacts with the Werner’s Syndrome Gene Product Physically and Functionally. J. Biol. Chem. 276, 20364–20369. 10.1074/jbc.C100035200.

56. Jurga, M., Abugable, A.A., Goldman, A.S.H., and El-Khamisy, S.F. (2021). USP11 controls R-loops by regulating senataxin proteostasis. Nat. Commun. 12. 10.1038/s41467-021-25459-w.

57. Murfuni, I., De Santis, A., Federico, M., Bignami, M., Pichierri, P., and Franchitto, A. (2012). Perturbed replication induced genome wide or at common fragile sites is differently managed in the absence of WRN. Carcinogenesis 33, 1655–1663. 10.1093/carcin/bgs206.

58. Kawabe, Y. ichi, Seki, M., Yoshimura, A., Nishino, K., Hayashi, T., Takeuchi, T., Iguchi, S., Kusa, Y., Ohtsuki, M., Tsuyama, T., et al. (2006). Analyses of the interaction of WRNIP1 with Werner syndrome protein (WRN) in vitro and in the cell. DNA Repair (Amst). 5, 816–828. 10.1016/j.dnarep.2006.04.006.

59. Socha, A., Yang, D., Bulsiewicz, A., Yaprianto, K., Kupculak, M., Liang, C.C., Hadjicharalambous, A., Wu, R., Gygi, S.P., and Cohn, M.A. (2020). WRNIP1 Is Recruited to DNA Interstrand Crosslinks and Promotes Repair. Cell Rep. 32, 107850. 10.1016/j.celrep.2020.107850.

60. Kusienicka, A., Czmoczek, A., and Gubała, J. (2021). Proximity Ligation Assay Detection of Protein – DNA Interactions — Is There a Link between Heme Oxygenase-1 and.

61. Huertas, P., and Aguilera, A. (2003). Cotranscriptionally formed DNA:RNA hybrids mediate transcription elongation impairment and transcription-associated recombination. Mol. Cell 12, 711–721. 10.1016/j.molcel.2003.08.010.

62. Boubakri, H., De Septenville, A.L., Viguera, E., and Michel, B. (2010). The helicases DinG, Rep and UvrD cooperate to promote replication across transcription units in vivo. EMBO J. 29, 145–157. 10.1038/emboj.2009.308.

63. Mirsanaye, A.S., Typas, D., and Mailand, N. (2021). Ubiquitylation at Stressed Replication Forks: Mechanisms and Functions. Trends Cell Biol. 31, 584–597. 10.1016/j.tcb.2021.01.008.

64. Li, X., and Manley, J.L. (2005). Inactivation of the SR protein splicing factor ASF/SF2 results in genomic instability. Cell 122, 365–378. 10.1016/j.cell.2005.06.008.

65. Wellinger, R.E., Prado, F., and Aguilera, A. (2006). Replication Fork Progression Is Impaired by Transcription in Hyperrecombinant Yeast Cells Lacking a Functional THO Complex. Mol. Cell. Biol. 26, 3327–3334. 10.1128/mcb.26.8.3327-3334.2006.

66. London, T.B.C., Barber, L.J., Mosedale, G., Kelly, G.P., Balasubramanian, S., Hickson, I.D., Boulton, S.J., and Hiom, K. (2008). FANCJ is a structure-specific DNA helicase associated with the maintenance of genomic G/C tracts. J. Biol. Chem. 283, 36132–36139. 10.1074/jbc.M808152200.

67. Sollier, J., Stork, C.T., García-Rubio, M.L., Paulsen, R.D., Aguilera, A., and Cimprich, K.A. (2014). Transcription-Coupled Nucleotide Excision Repair Factors Promote R-Loop-Induced Genome Instability. Mol. Cell 56, 777–785. 10.1016/j.molcel.2014.10.020.

68. Tuduri, S., Crabbé, L., Conti, C., Tourrière, H., Holtgreve-Grez, H., Jauch, A., Pantesco, V., De Vos, J., Thomas, A., Theillet, C., et al. (2009). Topoisomerase I suppresses genomic instability by preventing interference between replication and transcription. Nat. Cell Biol. 11, 1315– 1324. 10.1038/ncb1984.

69. Mohaghegh, P., Karow, J.K., Brosh, R.M., Bohr, V.A., and Hickson, I.D. (2001). The Bloom’s and Werner’s syndrome proteins are DNA structure-specific helicases. Nucleic Acids Res. 29, 2843–2849. 10.1093/nar/29.13.2843.

70. Paulsen, R.D., Soni, D. V., Wollman, R., Hahn, A.T., Yee, M.C., Guan, A., Hesley, J.A., Miller, S.C., Cromwell, E.F., Solow-Cordero, D.E., et al. (2009). A Genome-wide siRNA Screen Reveals Diverse Cellular Processes and Pathways that Mediate Genome Stability. Mol. Cell 35, 228–239. 10.1016/j.molcel.2009.06.021.

71. Bochman, M.L., Paeschke, K., and Zakian, V.A. (2012). DNA secondary structures: Stability and function of G-quadruplex structures. Nat. Rev. Genet. 13, 770–780. 10.1038/nrg3296.

72. Tsurimoto, T., Shinozaki, A., Yano, M., Seki, M., and Enomoto, T. (2005). Human Werner helicase interacting protein 1 (WRNIP1) functions as novel modulator for DNA polymerase δ. Genes to Cells 10, 13–22. 10.1111/j.1365-2443.2004.00812.x.

73. Rhodes, D., and Lipps, H.J. (2015). Survey and summary G-quadruplexes and their regulatory roles in biology. Nucleic Acids Res. 43, 8627–8637. 10.1093/nar/gkv862.

74. Varshney, D., Spiegel, J., Zyner, K., Tannahill, D., and Balasubramanian, S. (2020). The regulation and functions of DNA and RNA G-quadruplexes. Nat. Rev. Mol. Cell Biol. 21, 459– 474. 10.1038/s41580-020-0236-x.

75. Paeschke, K., Capra, J.A., and Zakian, V.A. (2011). DNA Replication through G-Quadruplex Motifs Is Promoted by the Saccharomyces cerevisiae Pif1 DNA Helicase. Cell 145, 678–691. 10.1016/j.cell.2011.04.015.

