## Supplementary Materials, Figure Legends and Figures for "WRNIP1 PREVENTS G4/R-LOOP-ASSOCIATED GENOMIC INSTABILITY"

### SUPPLEMENTARY MATERIALS AND METHODS

#### Site-directed mutagenesis and cloning

Site-directed mutagenesis of the WRNIP1 full-lenght cDNA (Open Biosystems) was performed on the pCMV-FLAGWRNIP1 plasmid that contains the wild-type ORF sequence of WRNIP1. Substitution of Asp37 to Ala in pCMV-FLAGWRNIP1 was introduced by the Quick-change XL kit (Stratagene) using mutagenic primer pairs designed, according to the manufacturer's instructions. Each mutated plasmid was verified by full sequencing of the WRNIP1 ORF.

#### Plasmids and RNA interference

The construct used to perform RNaseH1 overexpression experiments is a generous gift from Prof. R.J. Crouch (National Institutes of Health, Bethesda, USA). As previously described[1] , the GFP-tagged RNaseH1 plasmid was generated by introducing a mutation on Met27 abrogating mitochondrial localization signal (RNaseH1-M27). To express the plasmids, cells were transfected using the Neon™ Transfection System Kit (Invitrogen), according to the manufacturer's instructions. WRNIP1, FANCJ and RAD18 genetic knockdown experiments were performed by Interferin (Polyplus), according to the manufacturer's instructions. siRNAs were used at 20 nM for WRNIP1, 7,5 nM for FANCJ and 10 nM for RAD18. WRNIP1 depletion was achieved using a siRNA (QIAGEN) targeting the 3'UTR region of human protein (5'-ATGAATTAATGTTATAAGG-3'). Knockdown of FANCJ was obtained by transfection of siRNA Smartpool (Dharmacon, L-010587-00). Knockdown of FANCJ was obtained by transfection of Silencer Select siRNA by Thermo Fischer Scientific (s32297). As a control, a siRNA duplex directed against GFP was used. Depletion of the proteins was confirmed by Western blot using an anti-WRNIP1 antibody (Bethyl Laboratories), anti-FANCJ (Protein Tech) and anti-RAD18 (Abcam).

#### Chemicals

Chemicals used were commercially obtained for the replication stress-inducing drug: aphidicolin (Aph, Sigma-Aldrich), hydroxyurea (HU, Sigma-Aldrich), the transcription elongation inhibitors: 5,6-dichloro-1-ß-D-ribofurosylbenzimidazole (DRB, Sigma-Aldrich) and Cordycepin (Cordy, Sigma-Aldrich), the G4-stabilizing ligand pyridostatin (PDS, Sigma-Aldrich) and the proteasome inhibitor (MG132; Sigma-Aldrich). The final concentrations of the drugs used were: 0.4 or 5 µM aphidicolin, 4 mM HU, 50 µM DRB, 50 µM Cordy, 2 or 5 µM PDS and 10 µM MG132. Stock solutions for all the chemicals were prepared in DMSO at a concentration of >1000-fold except for

PDS, which was diluted in high purity water. The final concentration of DMSO in the culture medium was always < 0.1%.

**Chromosomal aberrations**

Cells for metaphase preparations were collected according to standard procedure and as previously reported [2]. Cell suspension was dropped onto cold, wet slides to make chromosome preparations. The slides were air dried overnight, then for each condition of treatment, the number of breaks and gaps was observed on Giemsa-stained metaphases. For each time point, at least 50 chromosome metaphases were examined by two independent investigators, and chromosomal damage was scored at 100× magnification with an Olympus fluorescence microscope.

**Alkaline and neutral Comet assay**

DNA breakage induction was examined by alkaline Comet assay (single-cell gel electrophoresis) in denaturing conditions as described [3]. Cell DNA was stained with a fluorescent dye GelRed (Biotium) and examined at 40× magnification with an Olympus fluorescence microscope and examined at 40× magnification with an Olympus fluorescence microscope. Slides were analysed by a computerized image analysis system (CometScore, Tritek Corp). To assess the amount of DNA damage, computer-generated tail moment values (tail length × fraction of total DNA in the tail) were used. A minimum of 200 cells was analysed for each experimental point. Apoptotic cells (smaller comet head and extremely larger comet tail) were excluded from the analysis to avoid artificial enhancement of the tail moment.

The occurrence of DNA double-strand breaks was evaluated by neutral Comet assay as described [4]. Cell DNA was stained with a fluorescent dye GelRed (Biotium). Slides were analysed as described above.

**DNA fiber analysis**

Cells were pulse-labelled with 50 µM 5-chloro-2’-deoxyuridine (CldU) and 250 µM 5-iodo-2’-deoxyuridine (IdU) at specified times, with or without treatment as reported in the experimental schemes. DNA fibres were prepared and spread out as previously reported [5]. For immunodetection of labelled tracks the following primary antibodies were used: anti-CldU (rat-monoclonal anti-BrdU/CldU; BU1/75 ICR1 Abcam, 1:100) and anti-IdU (mouse-monoclonal anti-BrdU/IdU; clone b44 Becton Dickinson, 1:10). The secondary antibodies were goat anti-mouse Alexa Fluor 488 or goat anti-rabbit Alexa Fluor 594 (Molecular Probes, 1:200). The incubation with antibodies was accomplished in a humidified chamber for 1 h at RT.

Images were acquired randomly from fields with untangled fibres using Eclipse 80i Nikon Fluorescence Microscope, equipped with a Video Confocal (ViCo) system. The length of green labelled tracks were measured using the Image-J software, and values were converted into kilobases using the conversion factor 1μm = 2.59 kb as reported [6]. A minimum of 100 individual fibres were analysed for each experiment and the mean of at least three independent experiments presented. Statistics were calculated using Graph Pad Prism Software.

#### *In situ* PLA assay

The *in situ* proximity-ligation assay (PLA; Sigma-Aldrich) was performed according to the manufacturer's instructions. Exponential growing cells were seeded into 8-well chamber slides (Lab-Tek, Sigma Aldrich) at a density of 1.5-2.5×$10^4$ cells/well. After the indicated treatment, cells were permeabilized with 0.5% Triton X-100 for 10 min at 4°C, fixed with 3% formaldehyde/ 2% sucrose solution for 10 min, and then blocked in 3% BSA/PBS for 15 min. After washing with PBS, cells were incubated with the two relevant primary antibodies. Antibody staining was carried out in the standard immunofluorescence procedure. The primary antibodies used were: anti-FLAG (mouse-monoclonal; Sigma-Aldrich, 1:250), anti-WRNIP1 (rabbit-polyclonal; Bethyl 1:500), anti-S9.6 (mouse-monoclonal; Kerafast, 1:100), anti-PCNA (rabbit-polyclonal; Abcam 1:500), anti-RNA polII (mouse-monoclonal; Santa Cruz, 1:200), Anti-DNA G-quadruplex (G4) (mouse-monoclonal; Sigma-Aldrich, 1:100), anti-FANCJ (rabbit-polyclonal; Protein Tech, 1:250), anti-IdU (rabbit-polyclonal anti-BrdU/IdU; GeneTex, 1:100), recombinant anti-S9.6 (rabbit-monoclonal; Kerafast, 1:100). The negative control consisted of using only one primary antibody. Samples were incubated with secondary antibodies conjugated with PLA probes MINUS and PLUS: the PLA Probe anti-Mouse PLUS and anti-Rabbit Minus (Sigma-Aldrich). The incubation with all antibodies was accomplished in a humidified chamber for 1 h at 37°C. Next, the PLA probes MINUS and PLUS were ligated using their connecting oligonucleotides to produce a template for rolling-cycle amplification. During amplification, the products were hybridized with red fluorescence-labelled oligonucleotide. Samples were mounted in Prolong Gold antifade reagent with DAPI (blue).
Images were acquired randomly using Eclipse 80i Nikon Fluorescence Microscope, equipped with a Video Confocal (ViCo) system.

#### Cell fractionation, immunoprecipitation, and Western blot analysis

Chromatin fractionation and immunoprecipitation experiments were performed as previously described [6]. Analysis of the distribution of proteins in the chromatin fraction was carried out by a standard protocol of chromatin fractionation [7]. Cells (1.5×$10^7$) were harvested using a cell scraper, centrifuged (2 min, 1.300 × g, 4°C), and then pellet was washed twice with PBS (2 min, 1.300 × g, 4°C). Cell pellets were resuspended in buffer A (10 mM HEPES pH 7.9, 10 mM KCl, 1.5 mM $MgCl_2$, 0.34 M sucrose, 10% glycerol, 1 mM DTT, supplemented with protease inhibitor cocktail). Triton X-100 (0.1%) was added, and the cells were incubated for 5 min on ice. Nuclei were collected in pellet by centrifugation (4 min, 1.300 × g, 4°C). The supernatant was discarded, nuclei washed once in buffer A, and then lysed in buffer B (3 mM EDTA, 0.2 mM EGTA, 1 mM DTT, supplemented with protease inhibitor cocktail). Insoluble chromatin was collected by centrifugation (4 min, 1.700 × g, 4°C), washed once in buffer B, and centrifuged again under the same conditions. The final chromatin pellet was resuspended in 2× sample loading buffer (100 mM Tris/HCl pH 6.8, 100 mM DTT, 4% SDS, 0.2% bromophenol blue and 20% glycerol), sonicated on ice, and boiled for 10 min at 95°C, then subjected to Western blot as reported below.

Briefly, for immunoprecipitation (IP) experiments, exponential growing HEK293T cells were cultured overnight at a density of $2.5 \times 10^6$ per 100 mm Petri dish and treated or not as indicated. After treatment, cells were collected and centrifuged. The cell pellets were resuspended in lysis IP buffer (0.5% Triton X-100, 150 mM NaCl, 1 mM EGTA, 50 mM Tris/HCl pH 8.0), freshly supplemented with protease and phosphatase inhibitor cocktails (SERVA), 15U/mL benzonase (Sigma-Aldrich) and 1 mM $MgCl_2$. DNA was subsequently fragmented by passing the lysed suspension 5 to 10 times through a needle attached to a 2mL syringe and incubated 40 min on ice. After centrifugation, for each IP sample, lysate was incubated with 20 µl anti-FLAG M2 magnetic beads (Sigma-Aldrich) overnight at 4°C. The IP reaction was washed four times with the IP buffer, incubated in 2× sample loading buffer (100 mM Tris/HCl pH 6.8, 100 mM DTT, 4% SDS, 0.2% bromophenol blue and 20% glycerol) for 10 min at 95°C, then subjected to Western blot as described below.

The proteins were resolved on polyacrylamide gels and transferred onto nitrocellulose membrane using the Trans-Blot Turbo Transfer System (Bio-Rad). The membranes were blocked using 5% NFDM in TBST (50 mM Tris/HCl pH 8, 150 mM NaCl, 0.1% Tween-20), and incubated with primary antibody for 1 h at RT or overnight at 4°C. The primary antibodies used for WB were rabbit-polyclonal anti-WRNIP1 (Bethyl Laboratories, 1:2500), mouse-monoclonal anti-FLAG (Sigma-Aldrich, 1:1000), mouse-polyclonal anti-GAPDH (Millipore, 1:5000), rabbit-polyclonal anti-LAMIN B1 (Abcam, 1:30000), rabbit-polyclonal anti-FANCJ (Protein Tech, 1:1500), rabbit-polyclonal anti-WRN (Abcam, 1:1000), rabbit-polyclonal anti-RAD18 (Abcam, 1:2000).

The membranes were incubated with horseradish peroxidase-conjugated goat specie-specific secondary antibodies (Santa Cruz Biotechnology, 1:20000), for 1 h at RT. Visualisation of the signal was accomplished using Western Bright ECL HRP substrate (Advansta) and imaged using Chemidoc XSR+ (Chemidoc Imaging Systems, Bio-Rad). Quantification was performed on scanned images of blots using ImageLab software (Bio-Rad), and values shown on the panels represent a fraction compared with the matched untreated control, normalized against the housekeeping proteinsimmunoblotting( Lamin B1 or GAPDH).

### REFERENCES

1. Cerritelli, S.M., Frolova, E.G., Feng, C., Grinberg, A., Love, P.E., and Crouch, R.J. (2003). Failure to produce mitochondrial DNA results in embryonic lethality in Rnaseh1 null mice. Mol. Cell 11, 807–815. 10.1016/S1097-2765(03)00088-1.

2. Pirzio, L.M., Pichierri, P., Bignami, M., and Franchitto, A. (2008). Werner syndrome helicase activity is essential in maintaining fragile site stability. J. Cell Biol. 180, 305–314. 10.1083/jcb.200705126.

3. Pichierri, P., Franchitto, A., Mosesso, P., Palitti, F., and Molecolare, C. (2001). Werner's Syndrome Protein Is Required for Correct Recovery after Replication Arrest and DNA Damage Induced in S-Phase of Cell Cycle. Mol. Biol. Cell 12, 2412–2421.

4. Murfuni, I., De Santis, A., Federico, M., Bignami, M., Pichierri, P., and Franchitto, A. (2012). Perturbed replication induced genome wide or at common fragile sites is differently managed in the absence of WRN. Carcinogenesis 33, 1655–1663. 10.1093/carcin/bgs206.

5. Leuzzi, G., Marabitti, V., Pichierri, P., and Franchitto, A. (2016). WRNIP 1 protects stalled forks from degradation and promotes fork restart after replication stress. EMBO J. 35, 1437–1451. 10.15252/embj.201593265.

6. Basile, G., Leuzzi, G., Pichierri, P., and Franchitto, A. (2014). Checkpoint-dependent and independent roles of the Werner syndrome protein in preserving genome integrity in response to mild replication stress. Nucleic Acids Res. 42, 12628–12639. 10.1093/nar/gku1022.

7. Méndez, J., and Stillman, B. (2000). Chromatin association of human origin recognition complex, cdc6, and minichromosome maintenance proteins during the cell cycle: assembly of prereplication complexes in late mitosis. Mol. Cell. Biol. 20, 8602–8612.

### LEGENDS TO SUPPLEMENTARY FIGURES

**Figure S1**. **Loss of WRNIP1 or its UBZ domain leads to sensitivity to Aph-induced MRS**
Representative Giemsa-stained metaphases of shWRNIP1$^{WT}$, shWRNIP1, shWRNIP1$^{D37A}$ and shWRNIP1$^{T294A}$ cells treated or not with 0.4 µM Aph for 24 h. Metaphases were collected by colcemid and fixed as reported in "Supplementary Materials and Methods". Arrows indicate chromosomal aberrations. Inset show enlarged metaphases for a better visualization of chromosomal aberrations.

**Figure S2**. **Transcription-dependent DNA damage in WRNIP1-deficient or WRNIP1 UBZ mutant cells**
(**A**) Analysis of DNA damage accumulation evaluated by alkaline Comet assay. MRC5SV, shWRNIP1, shWRNIP1$^{D37A}$ and shWRNIP1$^{T294A}$ cells were treated or not with 0.4 µM Aph for 24 h and with the inhibition of RNA elongation Cordycepin (Cordy), as reported in the experimental scheme, then cells were collected and subjected to Comet assay. The graph shows data presented as mean tail moment ± SE from three independent experiments. Horizontal black lines represent the mean (***, $P < 0.001$; ****, $P < 0.0001$; two-tailed Student's t test).
(**B**) Levels of γ-H2AX signal in cells treated as in (A), then subjected to immunostaining with an anti-γ-H2AX. The graph shows data presented as percentage of γ-H2AX-positive cells. Representative images of nuclei showing the different number of γ-H2AX foci (green) per nucleus are reported. Nuclei were counterstained with DAPI.

**Figure S3**. **Transcription influences DNA replication in cells lacking WRNIP1 or its UBZ domain upon MRS**
Experimental scheme of dual labelling of DNA fibers in MRC5SV, shWRNIP1 and shWRNIP1$^{D37A}$ cells under unperturbed conditions (**A**) or upon MRS (**B**). Cells were pre-treated with the inhibition of RNA elongation DRB, as reported in the experimental scheme, pulse-labelled with CldU, treated or not with 0.4 µM Aph, then subjected to a pulse-labelling with IdU. The graph shows the analysis of replication fork velocity (fork speed) in the cells. The length of the green tracks was measured. Mean values are represented as horizontal black lines (ns, not significant; **, $P < 0.01$; ***, $P < 0.001$; ****, $P < 0.0001$; two-tailed Student's t test). (**C** and **D**) The graphs show the percentage of red (CldU) tracts (stalled forks) or green (IdU) tracts (restarting forks) in the cells.

**Figure S4**. **PDS exacerbates accumulation of G4s in WRNIP1-deficient cells**

**(A)** Immunostaining with BG4 in MRC5SV, shWRNIP1, shWRNIP1$^{D37A}$ and shWRNIP1$^{T294A}$ cells treated or not with 5 µM PDS for 24 h. Visualization of DNA G4 structures was performed with the BG4 antibody as described in “Materials and Methods”. Immunostaining shows G4 foci (green) in the cells. Nuclei were counterstained with DAPI. Representative images are given. The graph shows the quantification of BG4 nuclear intensity per nucleus. Data are presented as means of three independent experiments. Horizontal black lines represent the mean ± SE (ns, not significant; * $P < 0.05$; ****, $P < 0.0001$; Mann-Whitney test).

**(B)** PLA assay to evaluate colocalization of WRNIP1 and G4 in MRC5SV cells treated or not with a siRNA for RAD18 and 0.4 µM Aph for 24 h. Cells were fixed and stained with antibodies against WRNIP1 and G4. Representative images are given. Each red spot represents a single interaction between protein and DNA. Nuclei were counterstained with DAPI. Dot plot shows the number of PLA spots per nucleus. Data are presented as means of three independent experiments. Horizontal black lines represent the mean ± SE (****, $P < 0.0001$; Mann-Whitney test).

**(C)** Evaluation of levels of DNA G4 structures by immunofluorescence in shWRNIP1$^{WT}$ and shWRNIP1 cells pre-treated with Aph and DRB, and then exposed or not to PDS as indicated in the experimental scheme. Visualization of G4 was carried out as in (A). Representative images are given. The graph shows the quantification of BG4 nuclear intensity per nucleus. Data are presented as means of three independent experiments. Horizontal black lines represent the mean ± SE (ns, not significant; * $P < 0.05$; ** $P < 0.01$; ****, $P < 0.0001$; Mann-Whitney test).

**Figure S5**. **PDS enhances R-loop accumulation in the absence of WRNIP1**

Evaluation of R-loop levels by immunofluorescence analysis in MRC5SV, shWRNIP1 and shWRNIP1$^{D37A}$ cells treated or not with 5 µM PDS for 6 h. Cells were fixed and stained with anti-RNA-DNA hybrid S9.6 monoclonal antibody. Representative images are given. Nuclei were counterstained with DAPI. The graph shows nuclear S9.6 fluorescence intensity. The line represents the median value. Data are presented as means of three independent experiments. Horizontal black lines represent the mean. Error bars represent standard error (****, $P < 0.0001$; Mann-Whitney test).

**Figure S6**. **R-loop-dependent DSB formation in PDS-treated WRNIP1-deficient and WRNIP1 UBZ mutant cells**

(**B**) Analysis of DNA double-strand break (DSB) formation by neutral Comet assay. MRC5SV and shWRNIP1 cells were treated or not with 2 or 5 µM PDS for 24 h, and then subjected to Comet assay.

Dot blot shows data presented as mean tail moment ± SE from three independent experiments (ns, not significant; **** $P < 0.0001$; Mann-Whitney test). Representative images are given.

(**C**) Neutral Comet assay performed to evaluate the effect of transfection with GFP-tagged RNaseH1 on R-loop-mediated DSBs formation. Cells were treated or not with 2 or 5 µM PDS for 24 h after transfection with GFP-tagged RNaseH1 or empty vector. Dot plot shows data presented as mean tail moment ± SE from three independent experiments. Horizontal black lines represent the mean (ns, not significant; * $P < 0.05$; ** $P < 0.01$; **** $P < 0.0001$; Mann-Whitney test). Representative images are given.

**Figure S7. Transcription and replication inhibition prevents DSB formation in PDS-treated WRNIP1-deficient cells**

Analysis of DSB formation by neutral Comet assay in shWRNIP1$^{WT}$ and shWRNIP1 cells pre-treated with Aph and DRB, and then exposed or not to PDS as indicated in the experimental scheme. Dot plot shows data presented as mean tail moment ± SE from three independent experiments. Horizontal black lines represent the mean (ns, not significant; ** $P < 0.01$; *** $P < 0.001$; **** $P < 0.0001$; Mann-Whitney test). Representative images are given.

**Figure S8**. **Loss of WRNIP1 or its UBZ domain results in reduced levels of FANCJ**

(**A**) Analysis of chromatin binding of FANCJ in MRC5SV, shWRNIP1, shWRNIP1$^{D37A}$ and shWRNIP1$^{T294A}$ cells. Chromatin fraction of cells, treated or not with the 0.4 µM Aph for 24 h in combination or not with 50 µM DRB for 3 hours, were analysed by immunoblotting. The membrane was blotted with anti-FANCJ antibody. LAMIN B1 was used as a loading for the chromatin fraction. The amount of the chromatin-bound FANCJ is reported as ratio of FANCJ/LAMIN B1 normalized over the untreated control.

(**B**) Analysis of chromatin binding of FANCJ in MRC5SV and shWRNIP1 cells. Chromatin fraction of cells, cells pre-treated with Aph and DRB, and then exposed or not to PDS as indicated in the experimental scheme were analysed by immunoblotting, performed as in (A).

(**C**) Evaluation of FANCJ levels in U2OS cells upon MRS in combination or not with transcription inhibition. Whole cell extract (WCE) was analysed by immunoblotting. The membrane was blotted with anti-FANCJ antibody. GAPDH was used as a loading for the WCE. The amount of the FANCJ is reported as ratio of FANCJ/GAPDH normalized over the untreated control.

**Figure S9. Evaluation of the FANCJ depletion on DNA damage accumulation upon MRS**

Analysis of DNA damage levels by alkaline comet assay in MRC5SV and shWRNIP1 cells depleted or not of FANCJ. Cells were treated with 0.4 µM Aph for 24 h and 50 µM DRB for the last 3 hours and then subjected to Comet assay. The graph shows data presented as mean tail moment ± SE from three independent experiments. Horizontal black lines represent the mean (ns, not significant; ****, $P < 0.0001$; two-tailed Student's t test). Representative images are given.

**Figure S10. Effect of proteasome inhibition on co-localization of FANCJ and G4s upon MRS**

Analysis of PLA assay to evaluate co-localization of FANCJ and G4 in MRC5SV and shWRNIP1 cells treated or not with the proteasome inhibitor MG132. Cells were fixed and stained with antibodies against FANCJ and G4s. Representative images are given. Each red spot represents a single interaction between protein and DNA. Nuclei were counterstained with DAPI. Dot plot shows the number of PLA spots per nucleus. Data are presented as means of three independent experiments. Horizontal black lines represent the mean ± SE (ns, not significant; ****, $P < 0.0001$; Mann-Whitney test).

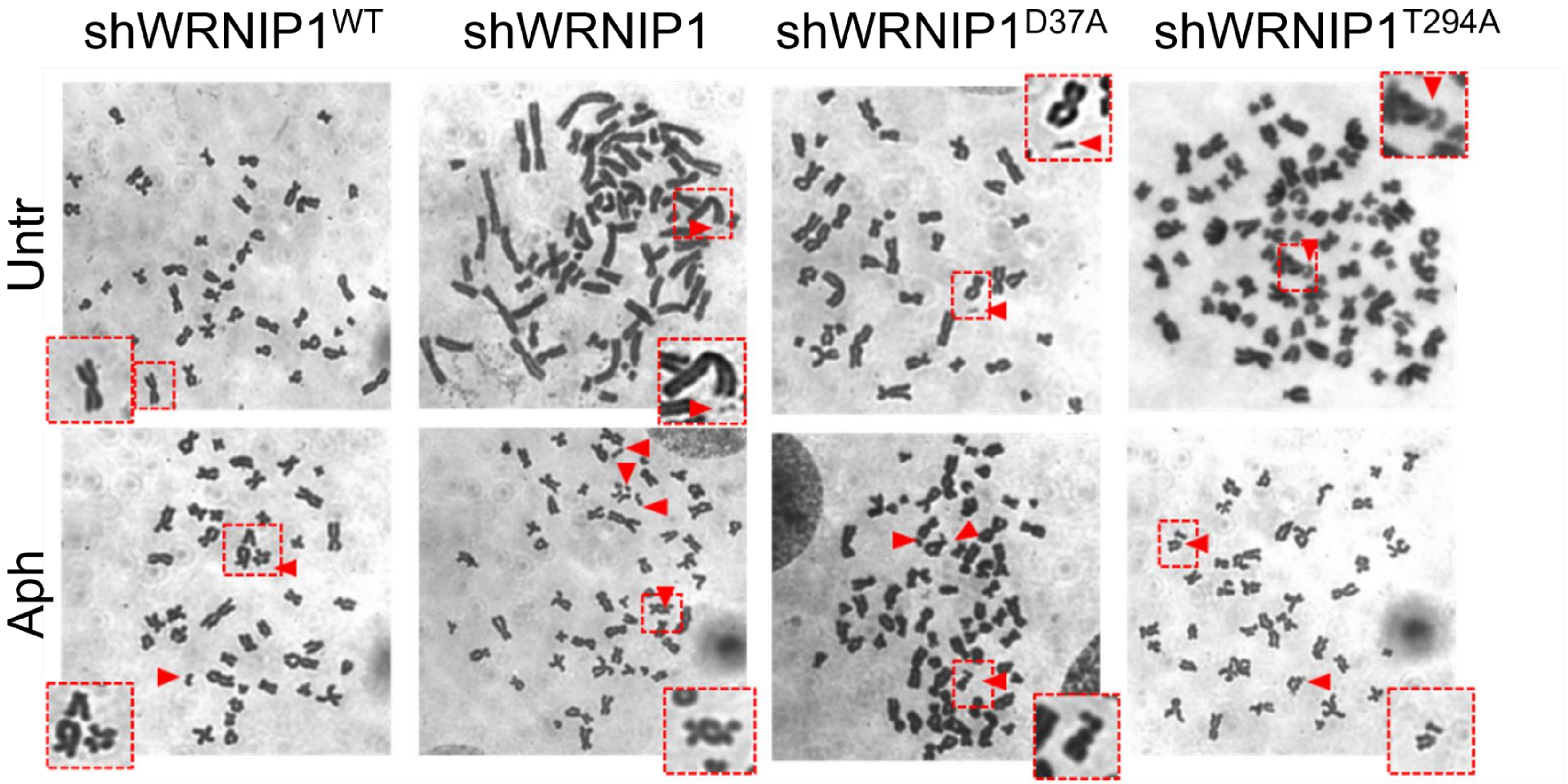


Suppl. Figure 1

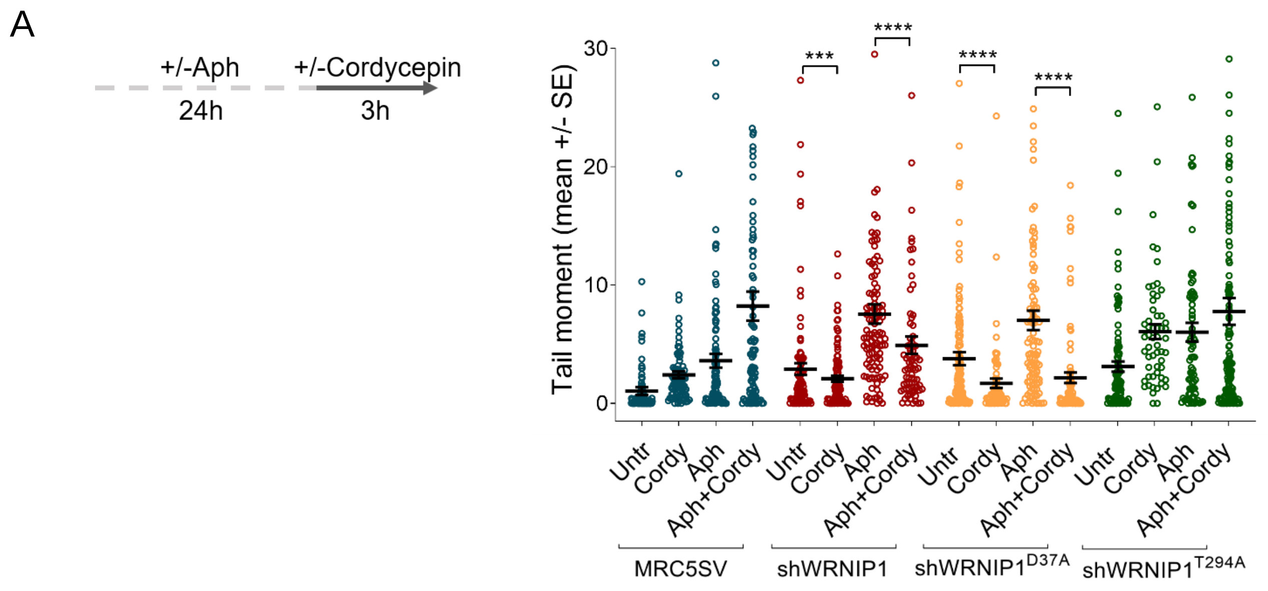


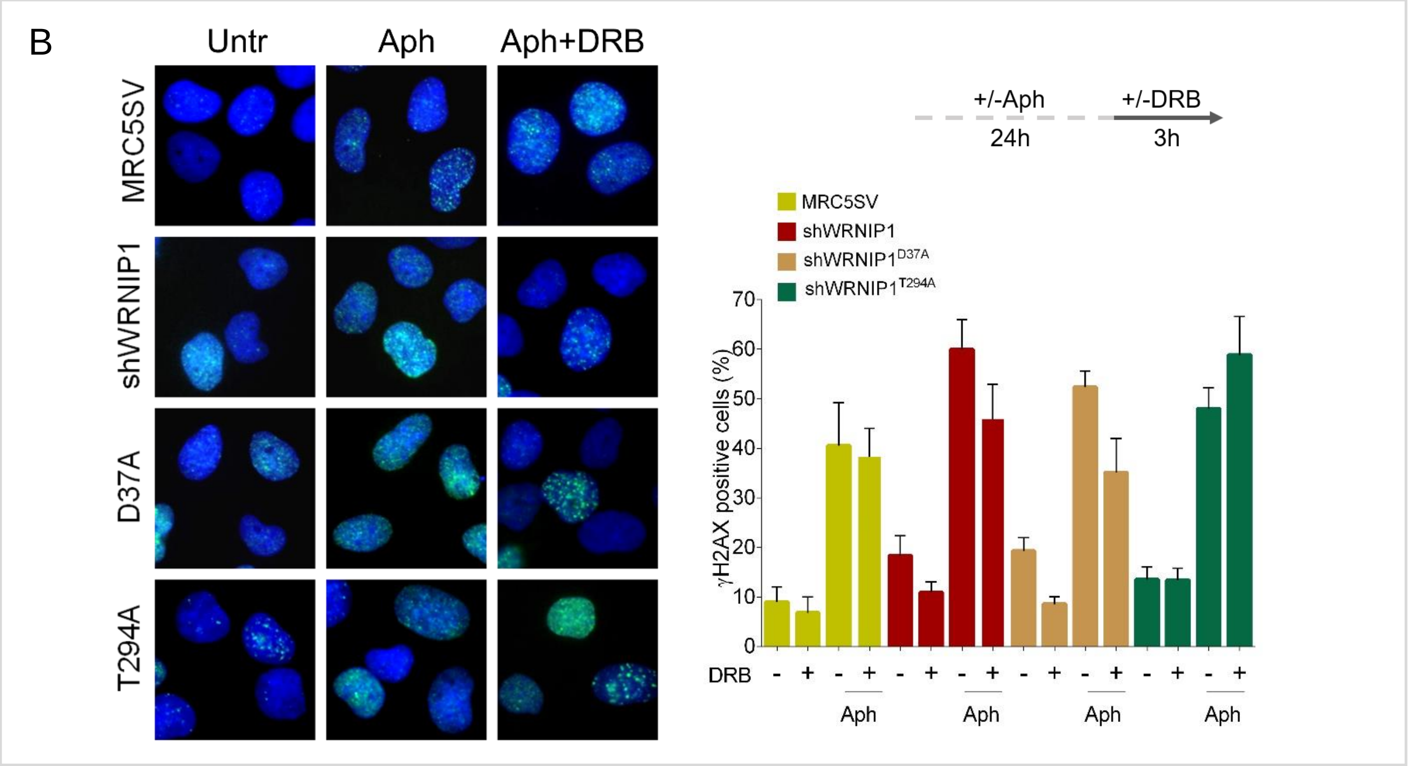


Suppl. Figure 2

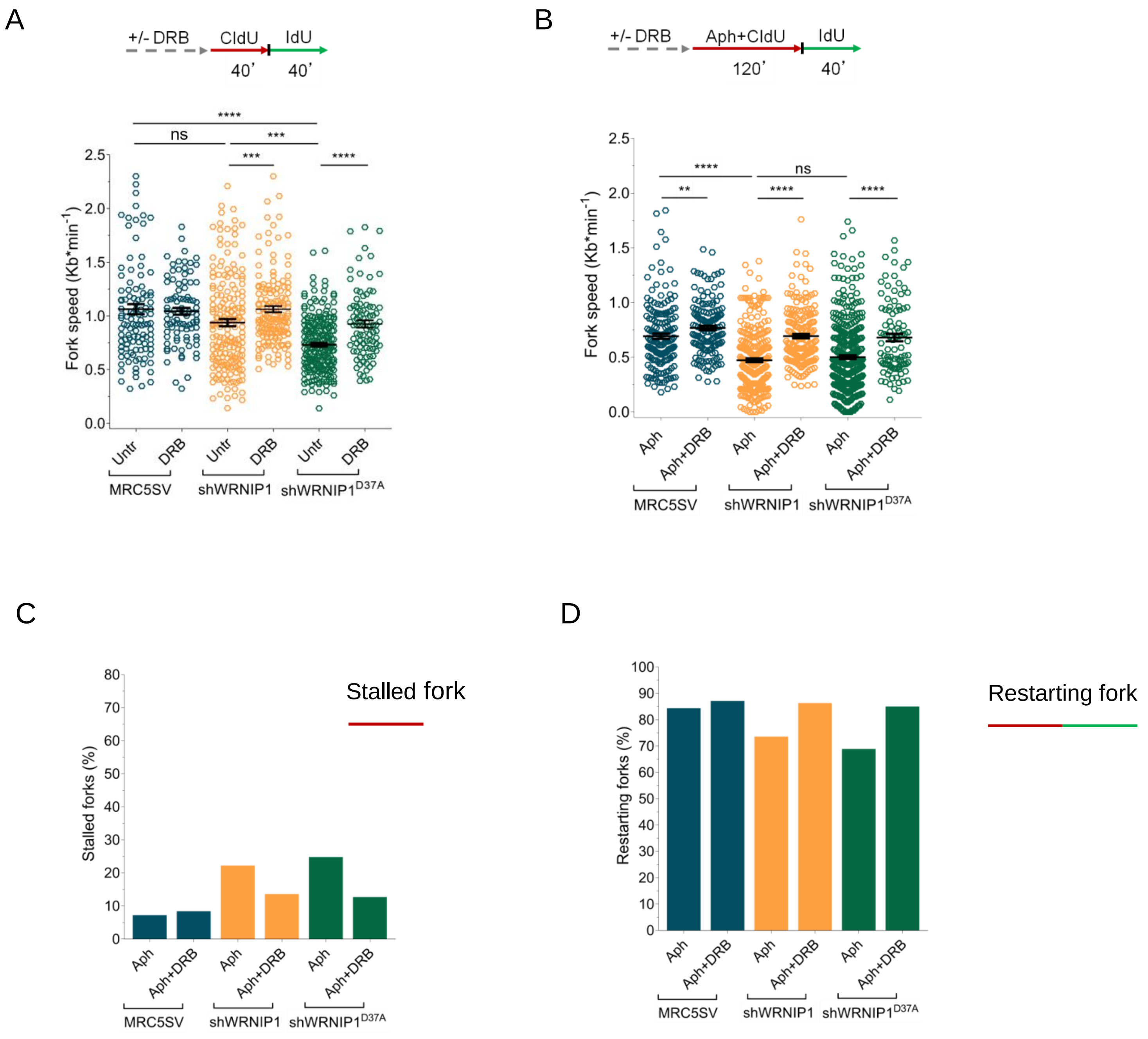


Suppl.Figure 3

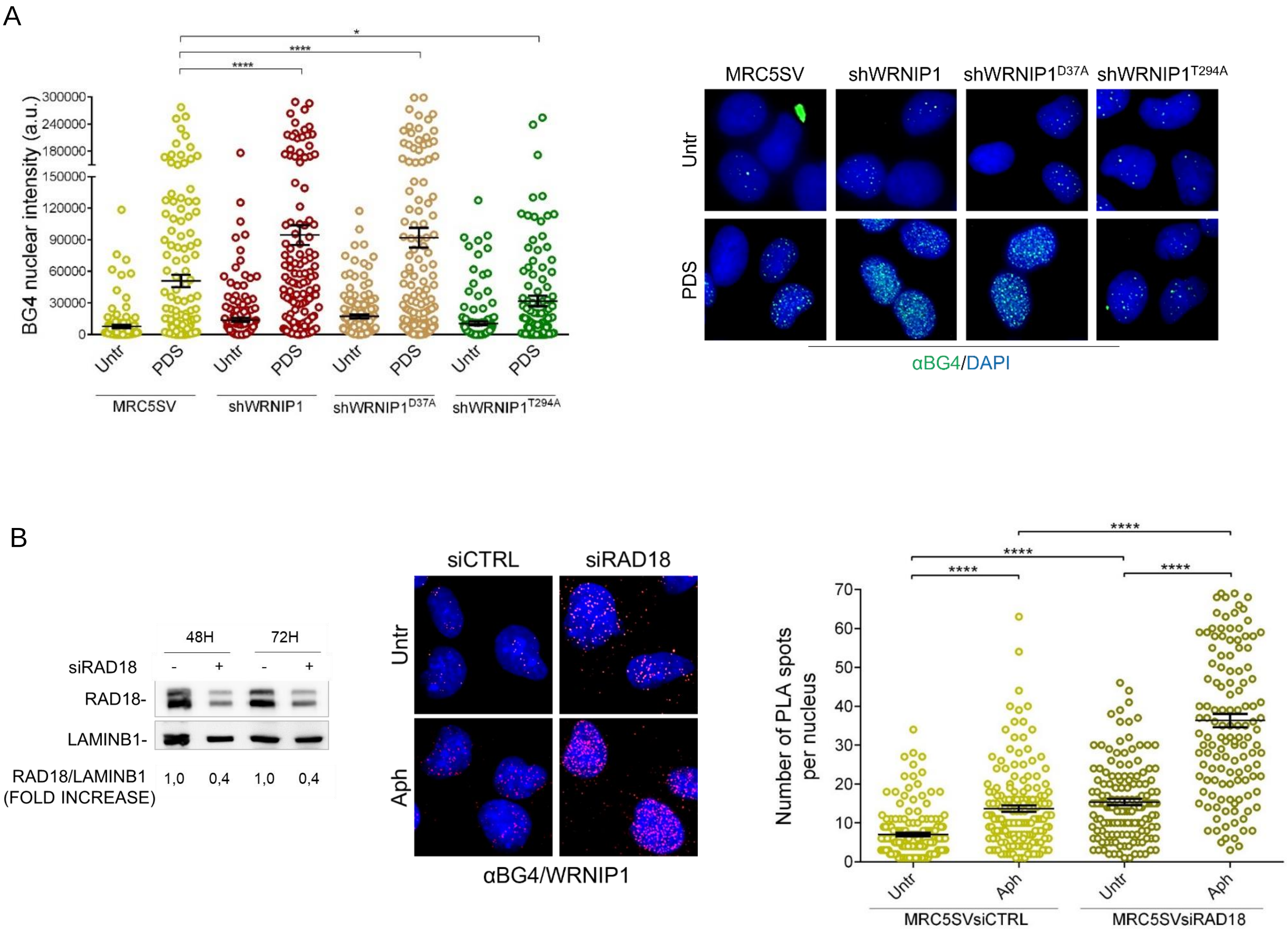


Suppl.Figure 4

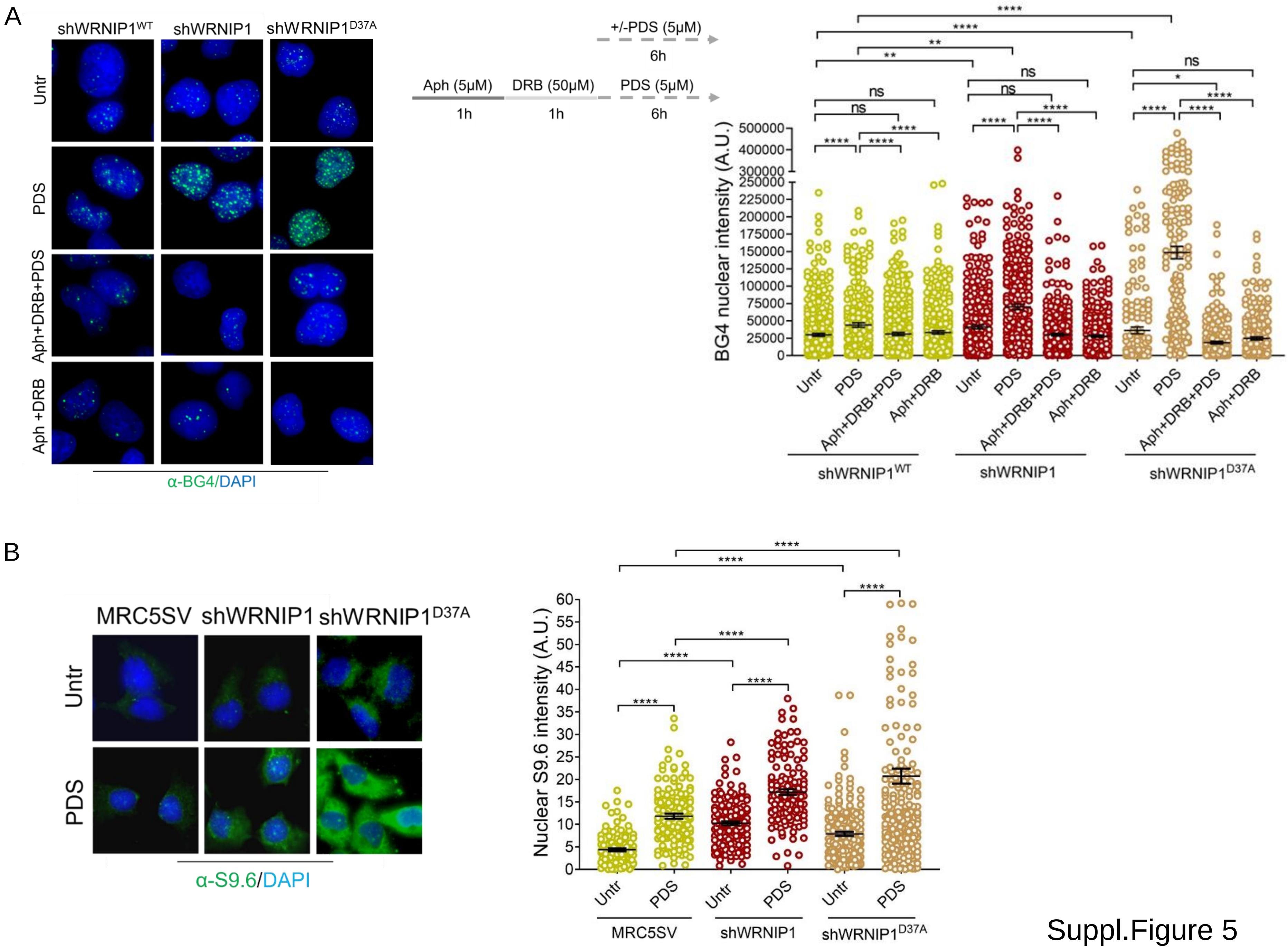


Suppl.Figure 5

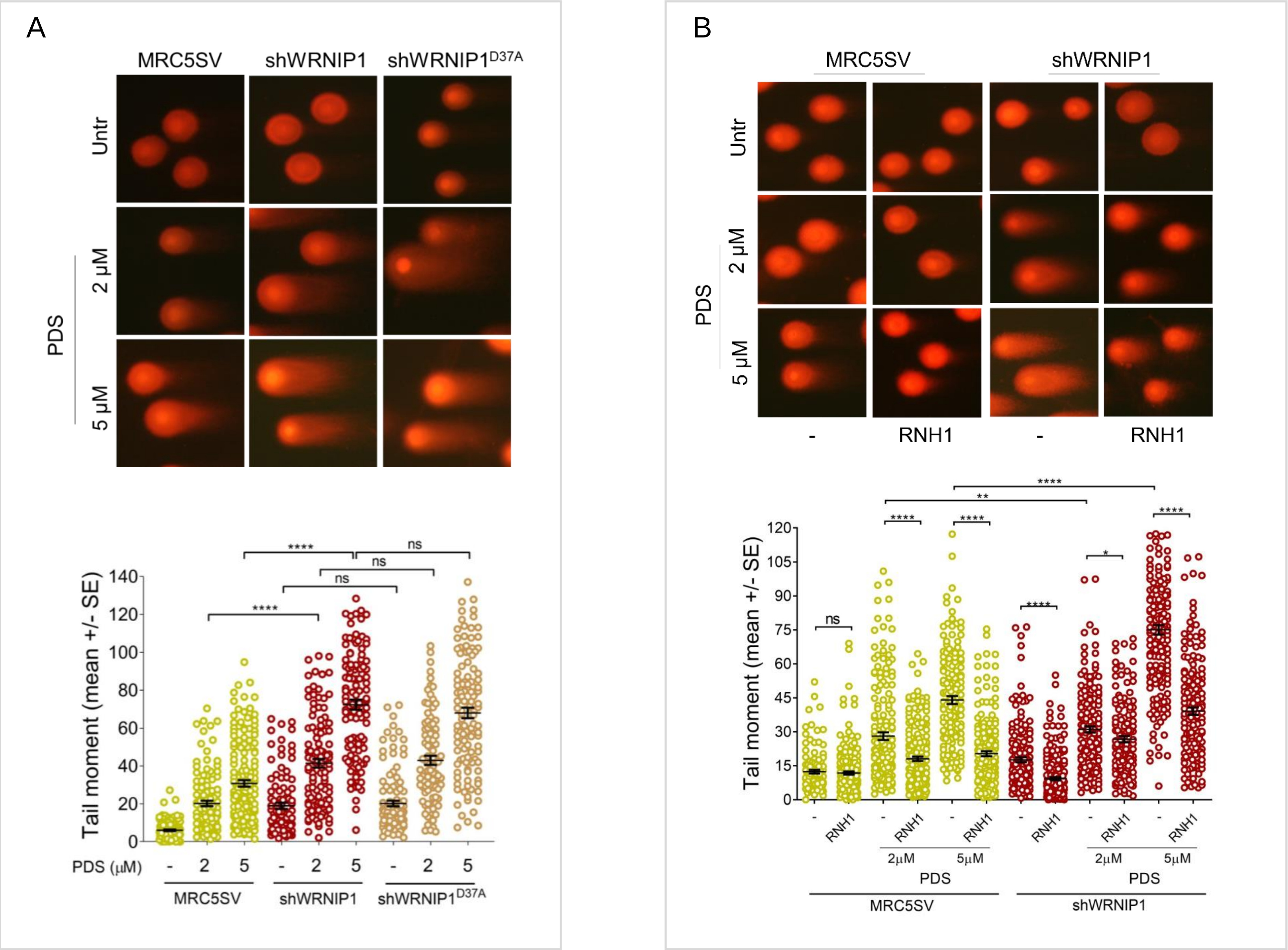


Suppl.Figure 6

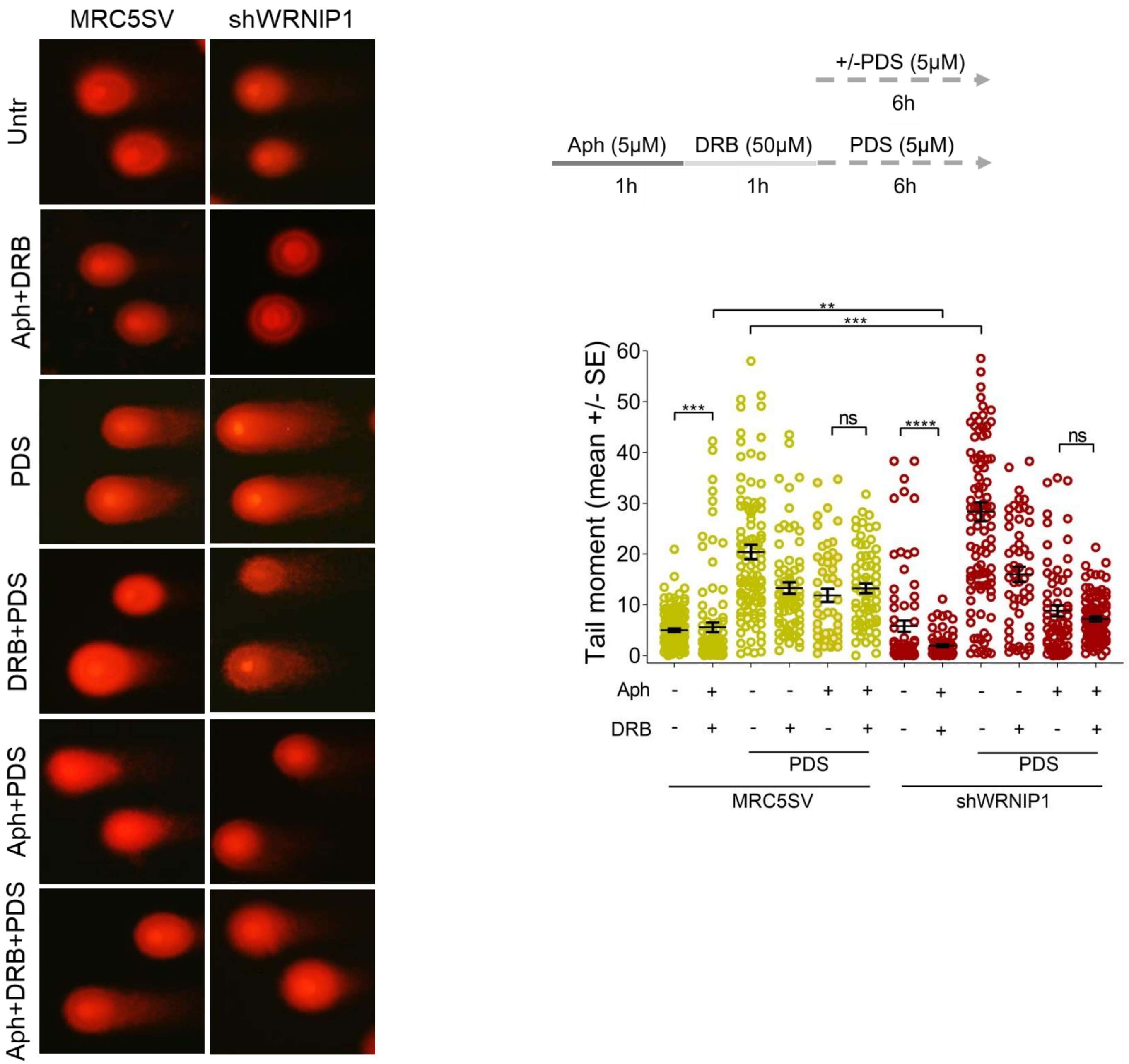


Suppl.Figure 7

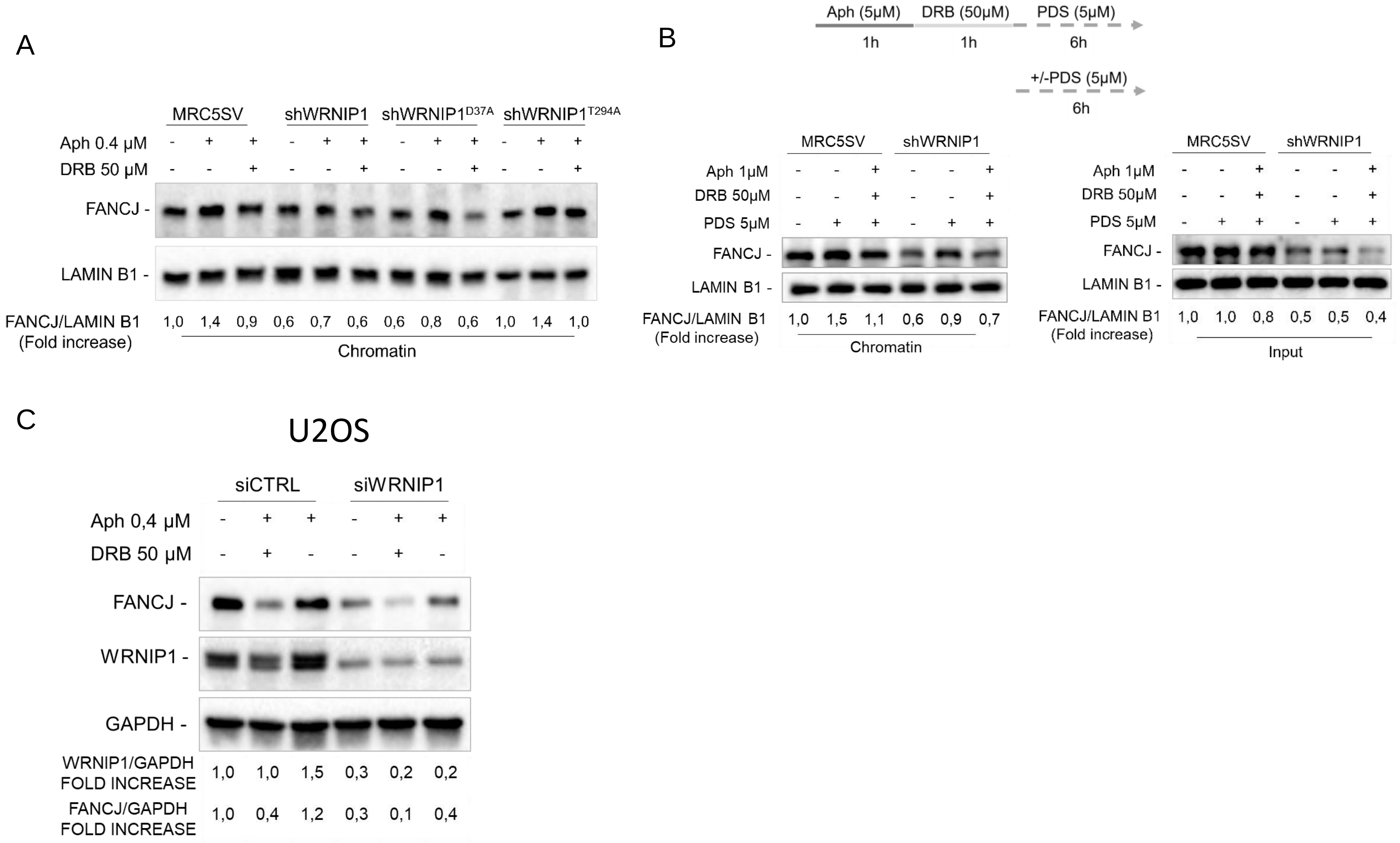


Suppl.Figure 8

A

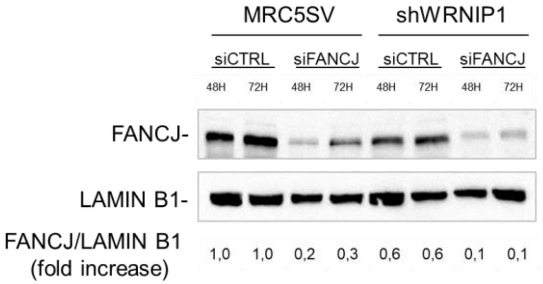


B

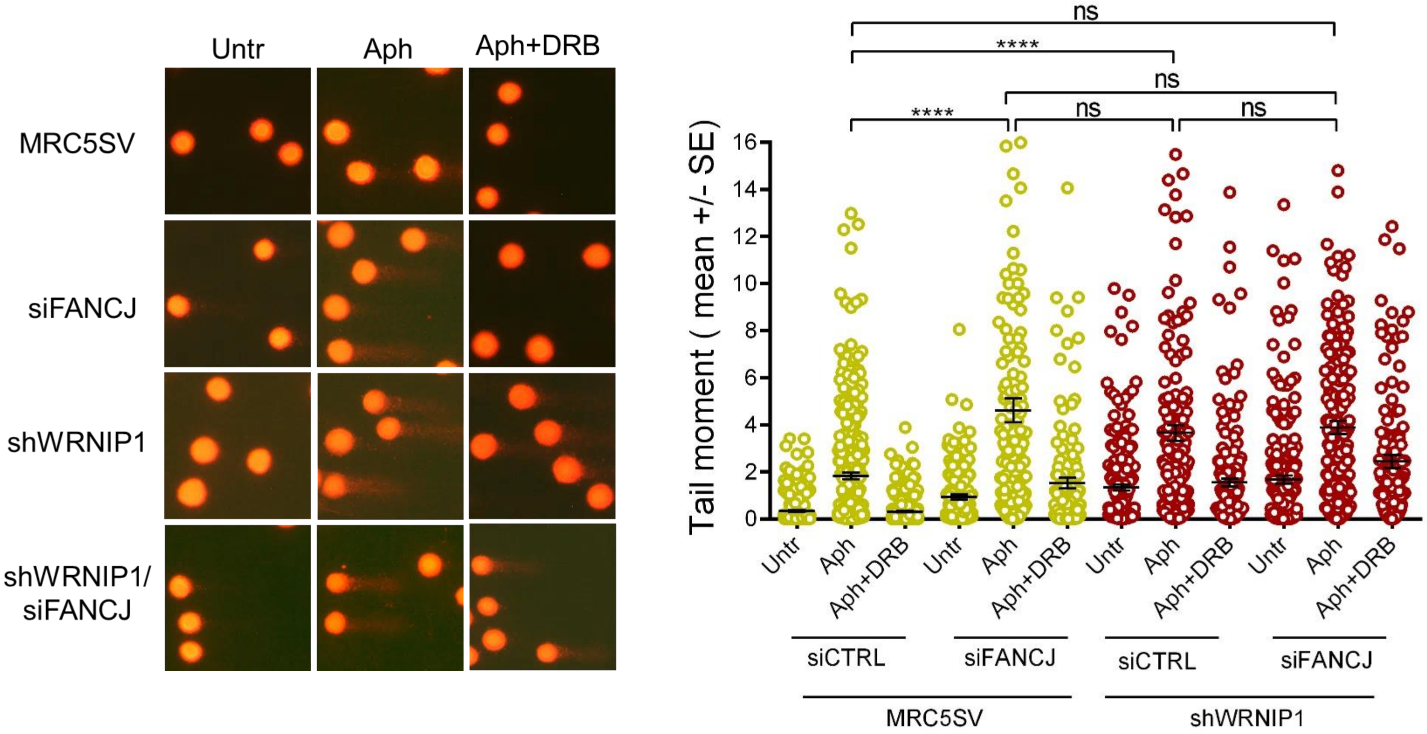


Suppl.Figure 9

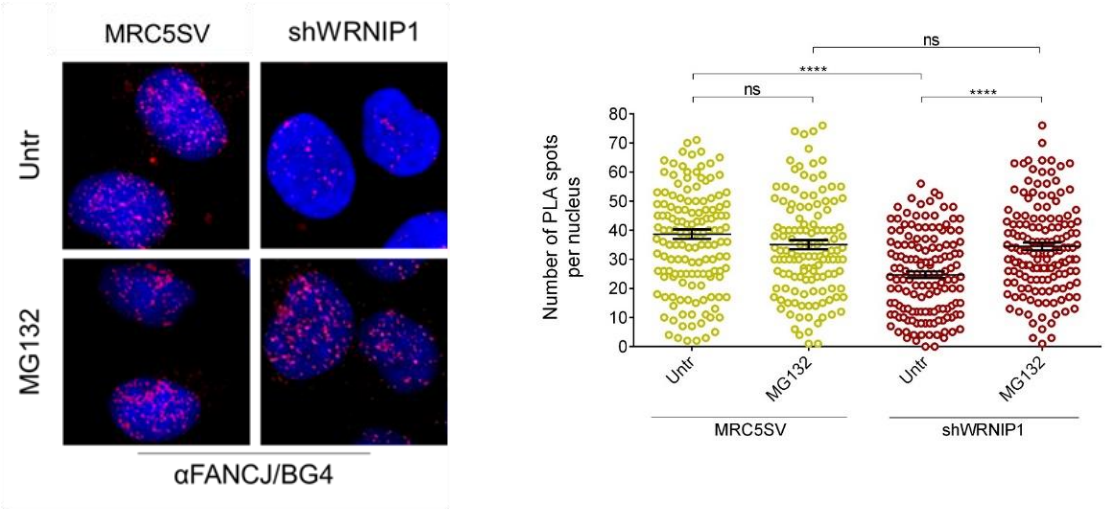


Suppl.Figure 10
